# Eubiota: Modular Agentic AI for Autonomous Discovery in the Gut Microbiome

**DOI:** 10.64898/2026.02.27.708412

**Authors:** Pan Lu, Yifan Gao, William G. Peng, Haoxiang Zhang, Kunlun Zhu, Elektra K. Robinson, Qixin Xu, Masakazu Kotaka, Harrison G. Zhang, Bingxuan Li, Anthony L. Shiver, Yejin Choi, Kerwyn Casey Huang, Justin L. Sonnenburg, James Zou

**Affiliations:** Department of Biomedical Data Science, Stanford University; Department of Computer Science, Stanford University; Department of Microbiology and Immunology, Stanford University; Biophysics Program, Stanford University; Department of Bioengineering, Stanford University; Institute for Human-Centered AI, Stanford University; Chan Zuckerberg Biohub, San Francisco

## Abstract

The gut microbiome regulates many aspects of human biology, including immunity and inflammatory diseases, yet mechanistic discovery and translation remain constrained by fragmented workflows and manual hypothesis integration. Here, we present **Eubiota**, an open-source modular agentic framework that decomposes microbiome inquiry using specialized agents for planning, execution, verification, and grounded synthesis. Coordinated through shared memory and domain-specific tools, **Eubiota** employs reinforcement learning to optimize multi-turn reasoning, achieving 87.7% benchmark accuracy and outperforming GPT-5.1 by 10.4%. Across four case studies, **Eubiota** enabled end-to-end discovery with experimental validation. Screening nearly 2,000 bacterial genes, **Eubiota** identified the *uvr-ruv* DNA repair axis as a fitness determinant under inflammatory stress, validated using transposon mutants and IBD metagenomes. It further designed a four-strain consortium that attenuated colitis severity in mice and generated a commensal-sparing antibiotic cocktail, demonstrating its utility in addressing community-level design challenges at cellular and molecular levels. Finally, **Eubiota** discovered diet-associated metabolites that suppress NF-*κ*B signaling. Together, these results establish **Eubiota** as a scalable, tool-grounded scientific copilot for mechanistically driven microbiome discovery.

## 1 Introduction

The human gut microbiome is a central regulator of host physiology [1, 2], impacting immune, metabolic, and neurobiological pathways that influence cancer, inflammatory disorders, and the gut–brain axis [3–6]. As its clinical relevance has grown, the field has shifted from descriptive correlations toward mechanistic frameworks aimed at elucidating how specific microbial genes, metabolites, and community structures exert causal effects on host physiology and disease progression [7–11]. Despite rapid data generation, microbiome discovery remains constrained by traditional paradigms [12, 13] in which data analysis, literature synthesis, hypothesis generation, and experimental design are performed in isolation and largely through manual effort [14, 15]. This fragmentation slows iteration, obscures mechanistic insight, and limits translation.

Artificial intelligence (AI) has emerged as a powerful catalyst for biological discovery, from protein structure prediction [16] to automated hypothesis generation [17, 18]. A new generation of agentic systems [19–25] has begun to assist experimental reasoning and workflow orchestration. However, many remain monolithic or rely on proprietary LLMs, limiting transparency, auditability, and domain-specific adaptation. Crucially, existing systems are often optimized for single tasks rather than the coordinated, iterative reasoning that microbiome discovery demands. In particular, the field lacks a cohesive framework that integrates heterogeneous models, tools, and domain constraints into a unified, end-to-end workflow tailored to microbiome and host health research [26–32].

The transition from data to biological discovery demands more than predictive accuracy; it requires hierarchical planning, adaptive tool use, and integration of heterogeneous evidence sources. High-level scientific objectives must be translated into reliable, auditable actions through iterative, stepwise adjustments that decouple execution from planning and are informed by intermediate results and feedback [33–37] across diverse resources, including the scientific literature, the open web, structured databases, and laboratory protocols, while respecting tool interfaces and domain constraints [38–40]. Because evidence is frequently incomplete, noisy, or conflicting, effective systems must incorporate verification mechanisms that assess consistency, determine when further analysis or evidence gathering is warranted, and terminate reasoning within practical time and computational budgets [41–43]. Finally, conclusions must be grounded in retrieved evidence and execution traces rather than generated as unconstrained narratives [44–46]. Together, these requirements motivate modular agentic architectures that decouple planning, execution, verification, and synthesis, enabling transparent, reliable integration of specialized tools and flexible operating modes. The result supports both interactive exploration and high-throughput discovery workflows in microbiome research, including the systematic exploration of combinatorial hypothesis spaces that are impractical to traverse by hand [21, 22, 47].

To address these challenges, we present **Eubiota**, a modular agentic AI framework for autonomous discovery in microbiome and host health research (Fig. 1**a**). Unlike other AI co-scientist systems that typically rely on closed models [20, 21, 25, 48], **Eubiota** supports open-weight backbones for all agents, enhancing transparency and extensibility. In this study, we optimized the central Planner using an open-weight model (Qwen3-8B) to enable trainable, auditable planning. **Eubiota** decomposes long-horizon scientific inquiry into four specialized agents for planning, execution, verification, and grounded generation, coordinated through a shared structured memory. This architecture enables integration of domain-specific models, databases, and analytical tools while maintaining transparency, auditability, and extensibility, and is complemented by an interactive interface that visualizes the reasoning process (Fig. 1**b**). To enhance reliability beyond standard orchestration, we introduce an agentic reinforcement learning strategy (GRPO-MAS) that optimizes multi-turn planning using outcome-based rewards. Across general biomedical and microbiome-specific benchmark evaluations, **Eubiota** (GRPO-MAS) achieves an average accuracy of 87.7%, outperforming general-purpose LLM baselines such as Qwen3-8B [49] (67.2%) and the proprietary GPT-5.1 [50] (77.3%), and surpassing a fixed Planner policy baseline (Frozen; 82.8% accuracy) (Fig. 1**c**).

**Figure 1:**
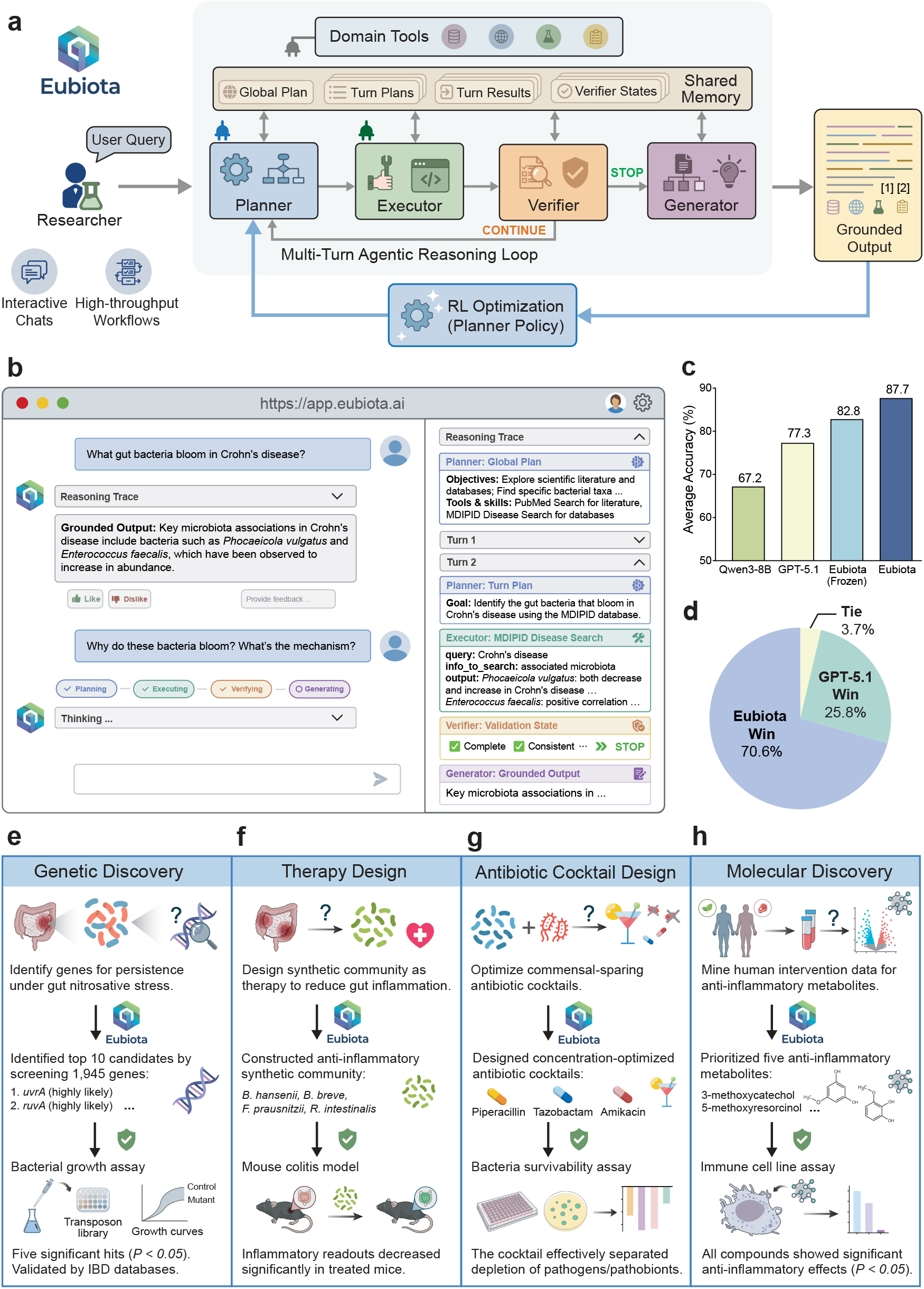
The Eubiota framework for autonomous microbiome discovery. **a**, Schematic of the **Eubiota** framework, comprising a multi-turn agentic reasoning loop orchestrated by specialized agents (Planner, Executor, Verifier, and Generator), shared Memory, domain tools, and outcome-driven reinforcement learning (RL) optimization. The Executor interfaces with external tools and records results in shared Memory; the Verifier evaluates evidence for consistency and sufficiency, determining whether to return control to the Planner (“CONTINUE”) or terminate (“STOP”). Upon termination, the Generator synthesizes a final response grounded in the full reasoning trace. The Planner policy is refined via the GRPO-MAS reinforcement learning strategy using trajectory-level outcome rewards to improve long-horizon planning. **b**, User interface for multi-turn queries, displaying a turn-by-turn reasoning trace and integrated user feedback. **c**, Average benchmark accuracy of **Eubiota** (87.7%) compared with a Frozen baseline (82.8%) and the general-purpose LLMs Qwen3-8B (67.2%) and GPT-5.1 (77.3%). **d**, Blinded domain-expert preference evaluation comparing **Eubiota** with the frontier model GPT-5.1. **e–h**, End-to-end biological discovery and design tasks. **e**, Identification of inflammatory stress resistance genes, including *uvrA* and *ruvA*. **f**, Design of a defined microbial consortium (*B. hansenii, B. breve, F. prausnitzii*, and *R. intestinalis*) for colitis. **g**, Optimization of a commensal-sparing antibiotic cocktail (piperacillin, tazobactam, and amikacin). **h**, Prioritization of anti-inflammatory metabolites (e.g., 3-methoxycatechol) from human intervention data.

To bridge the gap between computational reasoning and experimental practice, **Eubiota** operates in flexible modes spanning interactive querying (Fig. 1**b**) and large-scale, configuration-driven workflows that process thousands of candidates and connect directly to laboratory experimentation. In a blinded evaluation by domain scientists, **Eubiota** achieved a 70.6% win rate against GPT-5.1, with only 3.7% ties (Fig. 1**d**). An integrated interface enables human-in-the-loop collaboration through reasoning trace inspection and real-time corrective feedback. To ensure data sovereignty within clinical research settings, **Eubiota** supports full local deployment behind institutional firewalls. Together, these features allow researchers to steer inquiry while **Eubiota** scales evidence acquisition and iterative decision-making within explicit constraints.

We demonstrate the capabilities of **Eubiota** through four end-to-end discovery case studies spanning genes, communities, therapeutics, and metabolites. First, by integrating experimental phenotypes with genomic data, the system identified genes required for bacterial persistence under nitrosative stress, including *uvrA* and *ruvA*, which we validated using *Bifidobacterium breve* transposon mutants and confirmed to be enriched in human inflammatory bowel disease (IBD) cohorts (Fig. 1**e**). Second, **Eubiota** designed a defined microbial consortium that significantly attenuated inflammation in murine colitis (Fig. 1**f**). Third, it optimized a pathogen-biased antibiotic cocktail that preserved commensals (Fig. 1**g**). Finally, by mining dietary intervention data, **Eubiota** identified anti-inflammatory metabolites with effects validated in monocytic reporter assays (Fig. 1**h**). Across these applications, **Eubiota** transforms fragmented knowledge into auditable rationales and executable protocols, incorporating iterative feedback to refine proposals into experimentally testable designs. Rather than generating unconstrained narratives, these outputs provide a promising foundation for discovery at scale. Together, these results establish **Eubiota** as a scalable blueprint for collaborative, mechanistically grounded discovery in microbiome research.

## 2 System architecture and optimization

**Eubiota** employs a multi-agent architecture that decomposes scientific inquiry into four coordinated modules: planning, tool execution, verification, and grounded synthesis (Fig. 1**a**). Agents communicate through a structured shared memory that captures intermediate goals, tool interactions, and outputs, enabling transparent state tracking. By decoupling high-level reasoning from tool invocation and result auditing, the system facilitates the seamless integration of diverse domain resources while ensuring transparent and reproducible workflows. To enhance reliability in long-horizon tasks, the Planner is optimized via an outcome-driven reinforcement learning strategy that aligns multi-step reasoning with verifiable scientific outcomes.

### 2.1 Specialized agents and shared memory

Each scientific query initiates an iterative reasoning loop over discrete turns, as shown in Fig. 1**a**. The **Planner** translates the user query into an explicit multi-step strategy, identifying key scientific objectives, decomposing the task into ordered subgoals, and selecting appropriate tools based on their capabilities and constraints. The **Executor** interfaces directly with external tools; it executes the Planner’s intent by converting each planned action into validated commands. It enforces tool interface requirements (required fields, argument types, and allowable ranges), issues the call, and records the tool inputs and returned outputs in Memory. If generated parameters are invalid, the Executor automatically repairs formatting errors to match the tool’s documented fields and argument types.

In contrast to conventional planner-executor architectures [51–53], **Eubiota** incorporates a **Verifier** and a **Generator**. The Verifier audits accumulated evidence derived from tool outputs and retrieved citations stored in shared Memory. It evaluates completeness and sufficiency relative to the global plan and query, cross-checks claims against cited sources and prior-turn results, and identifies internal inconsistencies or unresolved ambiguities (for example, signaling “CONTINUE” when retrieval returns null results). The Planner uses this verification signal and rationale to refine the subsequent strategy, enabling iterative self-correction. Upon termination, the Generator synthesizes a final response grounded in the full execution trace.

A centralized **Memory** serves as the substrate for inter-agent communication, storing (i) the global plan, (ii) per-turn subgoals and tool selections, (iii) tool inputs and outputs, and (iv) verifier states. This shared state allows the Planner to adapt based on prior outcomes, minimizes redundant tool invocations, and enables the Verifier to audit the complete history. This persistent, structured record preserves context across agent transitions and ensures reproducible reporting of the evidence and reasoning underlying each conclusion.

### 2.2 Domain-specific toolset

To enable extensibility and reliability, **Eubiota** encapsulates each capability within a standardized Tool module defined by a rigorous schema, including typed inputs, output specifications, and usage constraints (Extended Data Fig. 1**a**). The Executor validates parameters against these schemas, employing an iterative self-correction mechanism to automatically repair malformed commands (Extended Data Fig. 1**b**). For knowledge-intensive tasks, database and document tools leverage retrieval-augmented generation (RAG) to synthesize context-aware evidence from embedded source records (Extended Data Fig. 1**c**). All interactions are logged in shared Memory, enabling the Verifier to audit intermediate evidence to ensure reproducible, traceable synthesis.

To support microbiome-specific reasoning, **Eubiota** integrates 18 tools spanning literature retrieval, biological databases, and laboratory resources (Extended Data Fig. 1**d**). Key capabilities include: (i) Grounded Search: Real-time web and literature retrieval via Google, Perplexity, Wikipedia, and PubMed to provide citation-backed evidence; (ii) Domain Databases: Structured queries to KEGG [54] and MDIPID [55] to map gene functions, drug interactions, and disease associations; (iii) Experimental Resources: Access to a curated Gene Phenotype Search (linking 118 gut bacteria to nitric oxide resistance) and a Protocol Search library of 55 assays; and (iv) Computational Utilities: A Python execution module (Python Coder) for quantitative analysis and context-aware document tools for user-provided data. The Planner dynamically selects from this toolkit based on the query and evolving Memory state.

For retrieval-intensive tools, source records are pre-indexed using a high-dimensional embedding model [56]. At runtime, the system retrieves the top-*k* semantically relevant matches (typically *k* = 10) and synthesizes them via an LLM to generate evidence-grounded responses. This approach allows the system to answer complex queries by drawing directly from verified source documents.

### 2.3 Iterative multi-turn orchestration

Reasoning unfolds iteratively over a maximum of *n*_max_ turns. At initialization, the Planner analyzes the query *q* and toolset 𝒯 to formulate a global strategy *p*_global_, which is persisted in Memory. Each turn *t* executes a four-stage cycle: (1) **Planning:** The Planner conditions on the current memory state ℳ_*t*_ to generate a turn-specific plan *p*_*t*_, specifying a tool *κ*_*t*_, a concrete subgoal, and required context; (2) **Execution:** The Executor translates *p*_*t*_ into a validated command *c*_*t*_, invoking the tool to yield result *r*_*t*_; (3) **Verification:** The Verifier audits the trajectory, returning a validation state *v*_*t*_ that determines whether to continue or terminate; and (4) **Update:** Memory is updated as ℳ_*t*+1_ = ℳ_*t*_ ∪ {*p*_*t*_, *c*_*t*_, *r*_*t*_, *v*_*t*_}. Let *T* ≤ *n*_max_ denote the terminal turn. Upon termination, the Generator synthesizes the final response grounded in the accumulated trace ℳ_*T*_.

Formally, we model this process as a Markov decision process (MDP) over states *S*_*t*_ = (*q*, ℳ_*t*_), where the query *q* is fixed and memory ℳ_*t*_ evolves. The Planner acts as a stochastic policy *π*_plan_(*p*_*t*_ | *S*_*t*_, 𝒯), selecting actions and tools *κ*_*t*_ ∈ 𝒯. Given plan *p*_*t*_, the Executor produces (*c*_*t*_, *r*_*t*_) and the Verifier returns *v*_*t*_. The state transition is defined as:

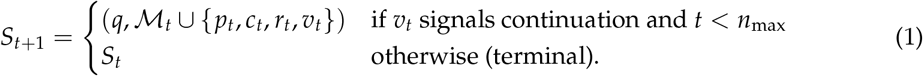

By conditioning on the full history ℳ_*t*_ embedded in *S*_*t*_, the system dynamically constructs adaptive tool sequences rather than following rigid, predefined paths. This modular architecture allows each agent to be instantiated with distinct LLM backbones, facilitating flexible, model-agnostic deployment. Formal inference procedures and prompt specifications are detailed in Supplementary Notes 2 and 4.

### 2.4 Optimization with agentic reinforcement learning

Off-the-shelf LLMs often struggle with the long-horizon reasoning required for scientific discovery, where early planning errors—such as querying irrelevant databases—can cascade into hallucinated conclusions. To mitigate this, we optimize the **Eubiota** Planner via a novel agentic reinforcement learning (RL) strategy that aligns multi-step reasoning with verifiable, trajectory-level outcomes (Extended Data Fig. 2**a**).

Unlike supervised fine-tuning, which enforces strict imitation of human demonstrations, our approach formulates planning as a learnable policy optimized via Group Relative Policy Optimization for Multi-Agent Systems (GRPO-MAS), adapted from the GRPO framework [57]. This strategy is critical for scientific inquiry, as it rewards any valid reasoning pathway that leads to the correct conclusion rather than mandating a single “gold standard” trajectory. During training, the system explores diverse multi-turn rollouts, receiving binary outcome rewards based on agreement with ground-truth labels (Extended Data Fig. 2**b**).

Crucially, GRPO-MAS propagates sparse terminal rewards back to intermediate planning steps, effectively assigning credit to the sequence of tool selections that produced a correct answer. Consequently, the Planner emergently learns robust scientific behaviors, such as cross-checking evidence, self-correcting upon failure, and prioritizing mechanistic data over superficial associations. This improves long-horizon reliability without requiring expensive step-by-step human annotation. To support this, we curated a heterogeneous training corpus of 2,000 instances dominated by microbiome reasoning (60%), but augmented with retrieval, general biomedical, and mathematical reasoning tasks (Extended Data Fig. 2**b**). We validate **Eubiota** (GRPO-MAS) across general and domain-specific benchmarks (Extended Data Fig. 2**c**; see Extended Methods).

### 2.5 Eubiota usage modes

To bridge the gap between exploratory reasoning and high-throughput discovery, **Eubiota** supports three distinct operating modes: (i) Interactive Chat for iterative, human-in-the-loop hypothesis generation; (ii) Batch Pipelines for large-scale, configuration-driven screening; and (iii) Local Deployment to ensure data sovereignty in clinical or proprietary settings. This flexibility allows researchers to seamlessly transition from initial inquiry to automated execution. Detailed specifications and implementation guides are provided in Supplementary Note 1.

## 3 Results

### 3.1 Benchmark evaluation of Eubiota

#### Evaluation setup

We assessed **Eubiota** across six multiple-choice benchmarks spanning general biomedical competence and microbiome-specific reasoning. For general biomedical knowledge, we evaluated performance on the Medicine subset of MedMCQA [58], which reflects professional clinical reasoning derived from medical entrance examinations, and WMDP-Bio [59], a benchmark probing expert-level biosecurity knowledge.

To evaluate microbiome-focused mechanistic reasoning, we constructed four domain-specific benchmarks derived from the microbiota-drug interaction and disease phenotype interrelation database (MDIPID) [55]. These tasks assess (i) drug impacts on microbial growth (Drug-Imp), mechanisms linking microbial activity to host proteins (MB-Mec), biological functions of proteins (Prot-Func), and mapping of expressed proteins to genes (Prot-Gen). Benchmark construction and evaluation protocols are detailed in Supplementary Note 5. To prevent data leakage, we removed near-duplicate training examples, standardized entity representations, and excluded any overlapping samples between training and evaluation sets.

Inference parameters were fixed to ensure reproducibility. Generation randomness was minimized (temperature = 0), with a maximum of 10 reasoning turns (*n*_max_ = 10) and a 300-second limit per query. Given the computational cost of multi-turn reasoning, we evaluated 100 randomly sampled instances per benchmark and report mean performance across three independent runs.

#### Performance analysis

Zero-shot base LLMs demonstrated limited accuracy on microbiome-specific benchmarks (Table 1). For example, the open-weights Qwen3-8B model [49] achieved 67.2% overall accuracy. Embedding the same backbone within **Eubiota**, with the Planner policy held fixed, increased accuracy to 82.8% (+15.6%), reflecting the contribution of domain-specific tools, structured memory, and iterative refinement through multi-turn reasoning. RL-optimized **Eubiota** (trained with the GRPO-MAS strategy) achieved the highest overall accuracy of **87.7%**. Gains were most pronounced on mechanistic reasoning tasks, with MB-Mec and Prot-Gen tasks reaching 94.0% and 94.7%, respectively. These improvements indicate that outcome-driven optimization enhances long-horizon planning and tool-use decisions, strengthening evidence acquisition and integration in microbiome-focused tasks. Encouraged by these results, we next applied **Eubiota** to four end-to-end experimental discovery settings.

**Table 1:** Performance on general biomedical and microbiome-specific benchmarks. We compare base LLMs (zero-shot) against the modular **Eubiota** framework (using a frozen Planner) and an RL-optimized variant (trained with GRPO-MAS). Accuracy scores are reported as mean ± s.d. across three independent runs. Δ denotes the percentage point improvement over the corresponding base model. Best results are highlighted in **bold**.

| Method | Planner setup |  | General biomedical |  | Microbiome-specific |  |  |  | Accuracy (%) |  |
| --- | --- | --- | --- | --- | --- | --- | --- | --- | --- | --- |
| | Backbone | Optimization | MedMCQA | WMDP-Bio | Drug-Imp | MB-Mec | Prot-Func | Prot-Gen | Avg. | $\Delta$ |
| *Base LLMs (zero-shot)* |  |  |  |  |  |  |  |  |  |  |
| Qwen3-8B [49] | — | — | 57.7±1.5 | 80.0±0.0 | 37.7±0.6 | 77.7±0.6 | 77.8±2.4 | 72.3±1.2 | 67.2 | — |
| GPT-4o [60] | — | — | 72.3±1.2 | 88.3±1.2 | 53.0±1.0 | 88.0±0.0 | 84.0±2.4 | 81.7±0.6 | 77.9 | — |
| GPT-5.1 [50] | — | — | **80.7±1.5** | **88.7±1.2** | 47.0±4.0 | 82.3±2.3 | 81.3±3.6 | 83.7±2.1 | 77.3 | — |
| *Ours (frozen Planner)* |  |  |  |  |  |  |  |  |  |  |
| **Eubiota** | Qwen3-8B | Frozen | 70.7±3.2 | 82.5±2.1 | 70.3±1.5 | 92.0±0.0 | **91.7±2.1** | 89.3±1.5 | 82.8 | ↑ 15.6 |
| **Eubiota** | GPT-4o | Frozen | 75.7±2.1 | 84.7±1.5 | **83.0±2.0** | 92.7±1.5 | 88.9±1.2 | 93.3±0.6 | 86.4 | ↑ 19.2 |
| **Eubiota** | GPT-5.1 | Frozen | 73.7±0.6 | **88.7±3.1** | 69.0±4.0 | 90.3±2.1 | 91.0±4.8 | 89.3±0.6 | 83.7 | ↑ 16.5 |
| *Ours (optimized Planner)* |  |  |  |  |  |  |  |  |  |  |
| **Eubiota** | Qwen3-8B | GRPO-MAS | 78.3±1.2 | 86.7±0.6 | 81.0±2.0 | **94.0±1.0** | **91.7±0.0** | **94.7±1.2** | **87.7** | ↑ 20.5 |

### 3.2 Eubiota driven inflammatory stress gene discovery

#### Nitrosative stress as a selective pressure in the inflamed gut

Chronic intestinal inflammation generates oxidative and nitrosative stress that reshapes gut microbial communities to dysbiosis by favoring stress-resilient taxa [61–64] (Fig. 2**a**). Among these pressures, host-derived nitric oxide (NO) functions as a key immune effector that perturbs bacterial physiology and contributes to inflammatory dysbiosis [65, 66]. We hypothesized that genes conferring resistance to nitrosative stress represent conserved fitness determinants in the inflamed gut.

**Figure 2:**
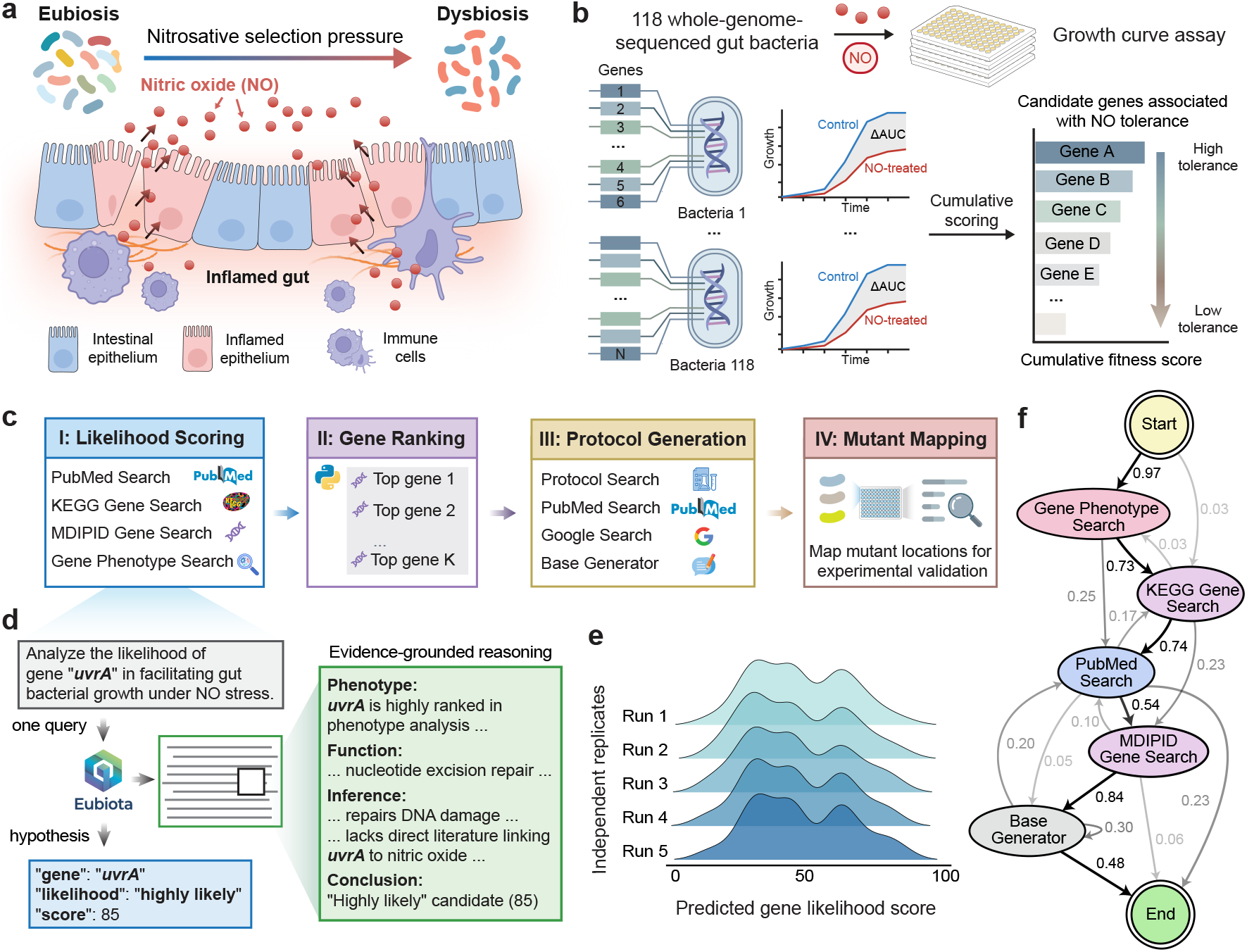
Eubiota-driven discovery of bacterial genes conferring resistance to inflammatory stress. **a**, Host-derived nitric oxide (NO) acts as a selection pressure in the inflamed gut, promoting the transition from eubiosis to dysbiosis. **b**, Genotype–phenotype association framework in which 118 whole-genome-sequenced gut bacterial strains were exposed to NO to quantify growth fitness (ΔAUC) and link phenotypic responses to gene content. **c**, Four-phase **Eubiota** workflow for prioritizing genes conferring nitrosative stress resistance: Phase I (Likelihood Scoring) integrates scientific literature, database queries, and phenotype data to assign likelihood scores; Phase II (Gene Ranking) selects top candidates; Phase III (Protocol Generation) produces literature-grounded validation protocols; and Phase IV (Mutant Mapping) identifies corresponding strains in an arrayed transposon mutant library. **d**, Representative reasoning trace for *uvrA*, illustrating how **Eubiota** links biological functions (nucleotide excision repair) to NO-stress phenotypes to support a highconfidence prediction. **e**, Gene prioritization stability across five independent workflow executions, shown as likelihood score distributions. **f**, State transition graph of autonomous reasoning. Nodes represent tools that are conditionally invoked to gather evidence across experimental data, pathway databases, and literature searches. Directed edges denote sequential tool invocations, with edge weights corresponding to cumulative transition frequencies across runs.

Systematic identification of such genes is challenging due to the extensive genetic diversity of gut microbes [67] and the difficulty of integrating phenotypic data, mechanistic annotation, and literature evidence at scale [68]. Although general-purpose LLMs can assist in information synthesis, they lack the explicit architectural grounding necessary for navigating domain-specific constraints, limiting their reliability in high-dimensional biological reasoning [41, 44, 45].

#### Phenotype-informed gene screening with Eubiota prioritization

To establish a scalable discovery pipeline, we quantified bacterial growth under nitrosative stress to enable gene-level inference. We cultured 118 whole-genome-sequenced isolates representing dominant human gut taxa [69] in the presence of 10 *µ*M DETA NONOate (NO donor), recapitulating inflammatory NO exposure [62, 65]. Growth kinetics were modeled using the Baranyi–Roberts equation [70, 71], and strain-level fitness changes were quantified as ΔAUC relative to untreated controls. Linking each annotated gene ortholog (defined by NCBI RefSeq functional nomenclature) to the NO-dependent fitness phenotype of its originating strain yielded 1,945 candidate genes associated with nitrosative stress tolerance (Fig. 2**b**). **Eubiota** prioritized these candidates through a four-phase workflow comprising evidence-grounded scoring, candidate filtering, experimental protocol generation, and transposon mutant mapping (Fig. 2**c**). For each gene, the system produced a structured assessment including a likelihood category, a quantitative score, and a concise mechanistic rationale supported by retrieved evidence (Fig. 2**d**).

#### Robustness and scale of agentic knowledge synthesis

To validate **Eubiota** ‘s reliability, we assessed prioritization stability across a random subset of 200 candidate genes. Across five independent runs, genes were scored consistently (mean Pearson *r* = 0.59) and highly reliable when aggregated (ICC(2,5) = 0.93), indicating that gene scores reflect systematic signal rather than stochastic variability. Overlapping score distributions across runs further supported stability (Fig. 2**e**, Supplementary Fig. 4**a, b**). Tool-use trajectories followed a reproducible progression from phenotype analysis to KEGG Gene Search to PubMed Search to MDIPID Search (Fig. 2**f**; Extended Data Fig. 4**a**), demonstrating task-aligned evidence integration. **Eubiota** analyzed 10,867 papers across 5 runs (Extended Data Fig. 4**b**), corresponding to a 38-fold increase in evidence acquisition efficiency relative to manual curation benchmarks[48] (Extended Data Table 1), scaling the labor-intensive process of biological evidence synthesis and providing a systematic framework for accessible large-scale discovery.

#### Discovery of a DNA repair axis required for NO tolerance

**Eubiota**-guided gene prioritization converged on a small, mechanistically coherent gene set, successfully validating known stress-response genes such as *ahpC* [72] while also discovering novel candidates. Among the top 10 candidates that mapped to the available *Bifidobacterium breve* UCC2003 transposon library (top 0.5% of global predictions; Extended Data Fig. 3**a, b**), the DNA repair genes *uvrA* and *ruvA* emerged as central hits. Although *uvrA* has been linked to inflammatory modulation in *Lactobacillus reuteri* [73] and *ruvA* to oxidative damage repair in *Escherichia coli* [74, 75], their roles in fitness under gut-relevant nitrosative stress have not been previously defined. Loss-of-function validation using single-gene transposon mutants [76] was employed to test these predictions (Fig. 3**a**). The experimental setup was guided by **Eubiota** generated protocols (Supplementary Note 6). Five of 10 prioritized mutants exhibited significant NO-dependent fitness defects (*P* < 0.05), quantified as log_2_(AUC_NO_/AUC_control_) [77] (Fig. 3**b**). In contrast, randomly selected genes showed no enrichment for inhibition (Extended Data Fig. 3**c, d**). The *ruvA* mutant displayed prolonged lag phase followed by biphasic growth, consistent with impaired resolution of stalled replication forks, whereas *uvrA* loss reduced rate of growth and decreased biomass accumulation (Fig. 3**c**). These phenotypes implicate DNA repair via nucleotide excision and homologous recombination pathways as critical determinants of bacterial fitness under inflammatory nitrosative stress. Importantly, analysis of the 118-strain growth dataset shows that inter-strain variation in these genes stratifies fitness under inflammatory nitrosative stress.

**Figure 3:**
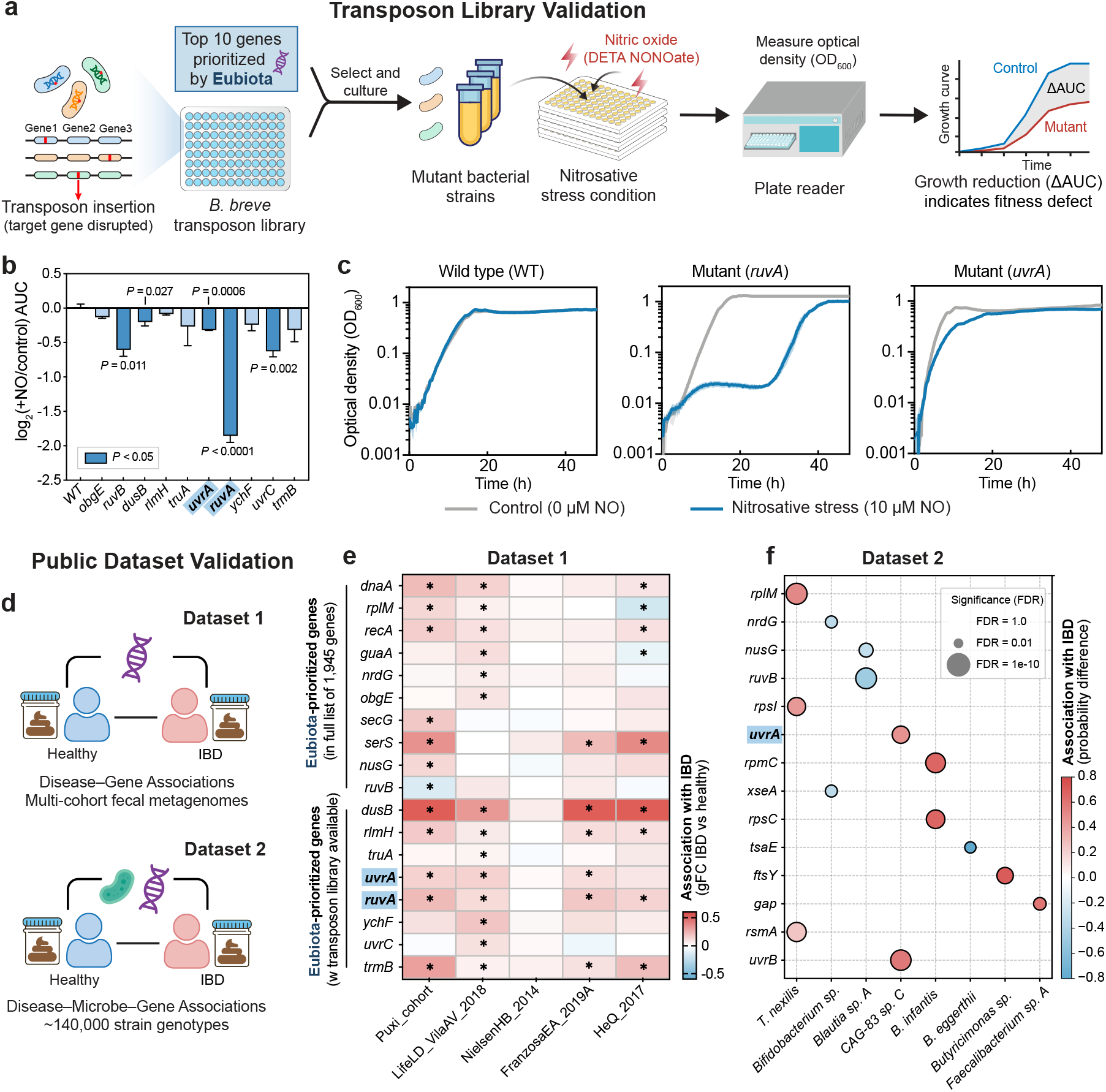
Experimental validation of Eubiota-prioritized bacterial genes. **a**, Validation strategy using a *B. breve* UCC2003 transposon library [76]. Selected loss-of-function mutants were exposed to nitric oxide (NO) generated by 10 *µ*M DETA NONOate. Fitness was quantified by comparing the growth area under the curve (AUC) between mutant and wild-type under NO treatment. **b**, Quantification of fitness effects across validated genes, shown as log_2_(AUC_NO_/AUC_control_) (mean ± s.e.m., *n* = 3 biological replicates). Statistical significance was determined by two-tailed Student’s *t*-test using AUC values. Mutants of *ruvB, dusB, uvrA, ruvA*, and *uvrC* exhibited significantly reduced growth (*P* < 0.05). **c**, Representative growth curves of selected transposon mutants under NO exposure compared with wild-type controls. Solid lines indicate the mean (*n* = 3); shading denotes s.e.m. **d**, Clinical relevance of **Eubiota**-prioritized genes assessed in large-scale human IBD cohorts. Candidate genes were evaluated using (i) multi-cohort fecal metagenome analyses of disease–gene associations (Dataset 1) [78] and (ii) strain-resolved disease–microbe–gene associations across ∼140,000 gut microbial genomes (Dataset 2) [79]. **e**, Heatmap of generalized fold change (gFC) values for **Eubiota**-prioritized genes across independent IBD cohorts [78]. Asterisks denote FDR-adjusted *P* < 0.05 within individual cohorts. **f**, Differential genetic probability between IBD-adapted and health-associated strains for genes ranked within the top 50 **Eubiota** predictions and present in the source dataset [79]. Dot size corresponds to FDR-adjusted *P* values reported in the original study.

#### Enrichment of stress-tolerance genes in human IBD metagenomes

To assess clinical relevance, we mapped prioritized genes to two independent public IBD metagenomic datasets (Fig. 3**d**). Cross-cohort analysis revealed consistent enrichment of DNA replication and repair genes, including *uvrA, ruvA, recA*, and *dnaA*, in patients with IBD relative to healthy controls (Fig. 3**e**) [78]. This pattern persisted at strain-level resolution across ∼140,000 reconstructed genomes from IBD and control metagenomes [79] (Fig. 3**f**). Notably, *uvrA/B* variants were significantly enriched within IBD-adapted *Oscillospiraceae* lineages. Beyond DNA repair, genes involved in translational capacity and fidelity (e.g., *rplM, rsmA*) were also enriched, suggesting coordinated adaptation to maintain genomic and proteostatic integrity under inflammatory stress. Together, these findings link experimentally validated nitrosative stress tolerance genes to *in vivo* selection in IBD, highlighting candidate fecal biomarkers for IBD diagnosis and high-priority targets for future mechanistic investigation and translational intervention.

### 3.3 Eubiota assisted therapeutic design for gut diseases

#### Rationale and challenges in therapeutic consortium design

Moving from single-gene discovery to community-level intervention, we next applied **Eubiota** to design synthetic microbial therapeutics. Defined consortia offer a controlled alternative to the heterogeneity of fecal microbiota transplantation (FMT) [80–82], yet selecting optimal species combinations requires balancing ecological stability, metabolic output, and host immune interactions [83–85] (Fig. 4**a**). General-purpose LLMs often provide unstable or poorly supported recommendations [32, 86], limiting their utility for experimental design. As an initial evaluation, we tested whether **Eubiota** could correctly prioritize a canonical consortium of *Roseburia intestinalis, Intestinimonas butyriciproducens, Faecalibacterium prausnitzii*, selected for their established production of the anti-inflammatory short-chain fatty acid (SCFA) butyrate [87–95]. While many factors in the microbiota can influence inflammation, the well-described impact of major fermentation end-products provided a high-confidence benchmark for a focused hypothesis. To control for positional bias, we permuted the order of candidate consortia across multiple trials. **Eubiota** consistently identified the butyrate-producing community regardless of its assigned label (A, B, or C) with comprehensive literature evidence and KEGG Organism Search, demonstrating that its selections are driven by biological reasoning rather than prompt structure (Extended Data Fig. 5**a, b**).

**Figure 4:**
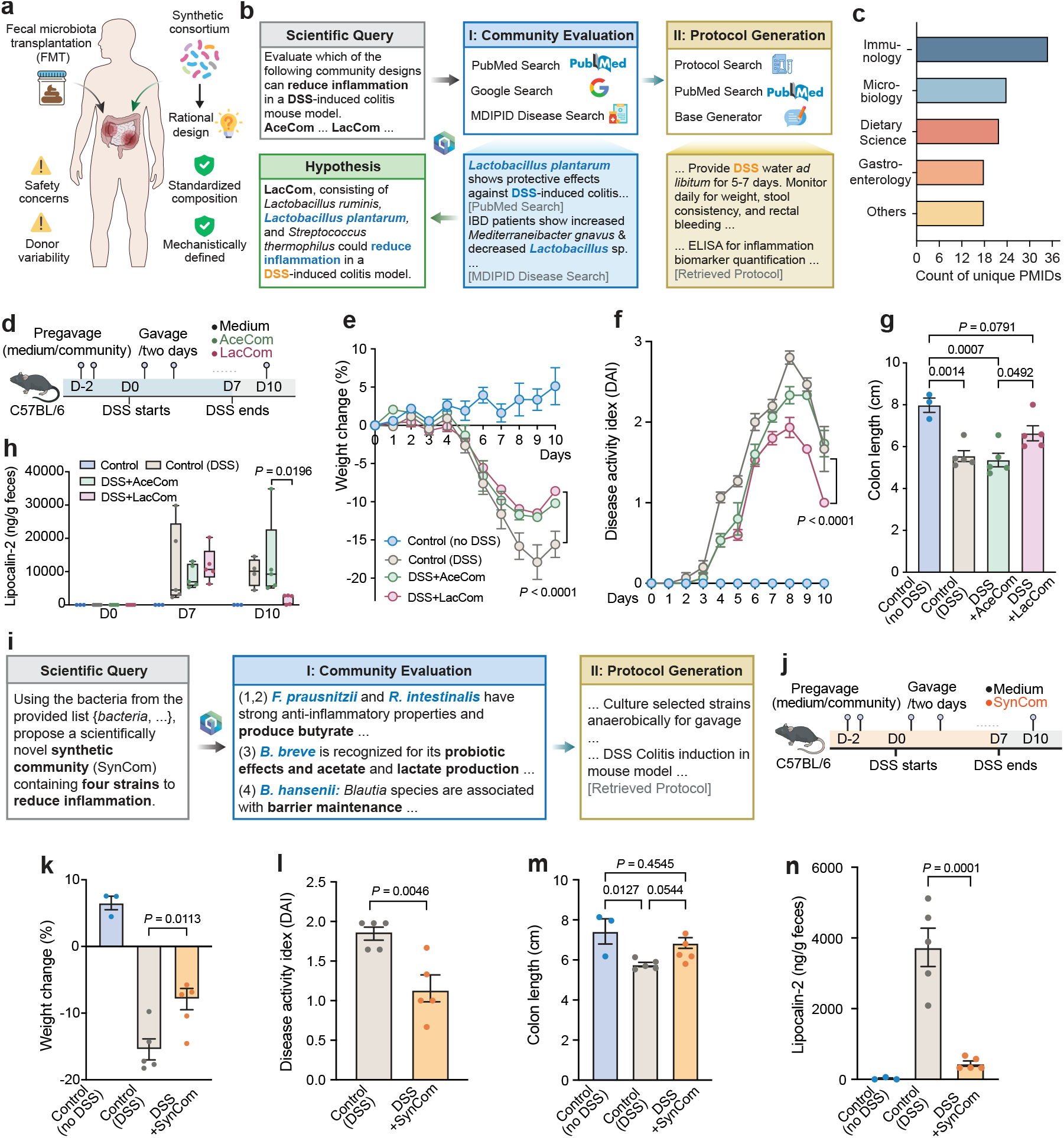
Eubiota-driven design and *in vivo* validation of synthetic microbial consortia. **a**, Conceptual framework for replacing fecal microbiota transplantation (FMT) with standardized, mechanistically defined synthetic microbial communities. **b**, Two-phase **Eubiota** workflow for therapeutic consortium design. Phase I integrates literature and database evidence to evaluate candidate communities and nominate an optimal design; Phase II generates a fully specified experimental protocol for validation in a DSS-induced colitis mouse model. **c**, Distribution of scientific literature analyzed during community evaluation, stratified by discipline and quantified by unique PubMed identifiers (PMIDs). **d**, Experimental design schematic for DSS-induced colitis validation following the **Eubiota**-generated protocol. **e–h**, *In vivo* comparison of DSS-treated mice receiving acetate-producing (AceCom) or lactate-producing (LacCom) consortia. During the recovery phase (days 7–10), LacCom-treated mice exhibited reduced weight loss (**e**) and lower Disease Activity Index (DAI) scores (**f**). At day 10, LacCom-treated mice showed increased colon length (**g**) and reduced fecal lipocalin-2 (LCN-2) levels (**h**) relative to DSS-only controls. **i**, Representative **Eubiota** reasoning trace for *de novo* consortium design (SynCom; *F. prausnitzii, R. intestinalis, B. breve*, and *B. hansenii*), highlighting prioritization based on predicted anti-inflammatory and butyrate-producing functions. **j**, Experimental design schematic for DSS-induced colitis validation of the SynCom consortium. **k–n**, Phenotypic outcomes of SynCom treatment in DSS-induced colitis at day 10. SynCom administration significantly reduced weight loss (**k**) and DAI scores (**l**), increased colon length (**m**), and decreased fecal LCN-2 levels (**n**) relative to DSS-only controls. Data are shown as mean ± s.e.m. (*n* = 3 for control; *n* = 5 per DSS and treatment group). Statistical significance was assessed by two-way ANOVA for **e, f, h**; one-way ANOVA with multiple-comparison correction for **g, k, m, n**; and two-tailed Student’s *t*-test for **l**.

We then asked **Eubiota** to evaluate two candidate communities for reducing DSS-induced colitis: an acetate-producing consortium (AceCom) and a lactate-producing (LacCom) consortium, addressing a persistent debate regarding the possible pro- versus anti-inflammatory roles of acetate and lactate in intestinal ecosystems [96–101]. In the absence of consensus on the superior metabolic strategy for attenuating gut inflammation, this setting provides a stringent test of **Eubiota** ‘s ability to adjudicate between competing, mechanistically complex intervention strategies (Fig. 4**b**), while remaining aware of the broader metabolic and immunological landscape that modulates these primary effects. **Eubiota** implemented a structured two-phase workflow.

In phase I, it synthesized mechanistic and empirical evidence from the literature and domain-specific databases to rank candidate consortia. In phase II, it translated the top-ranked hypothesis into an executable mouse study protocol to test the prediction, specifying dosing, controls, and readouts (Supplementary Note 6). Across repeated runs incorporating 90 relevant publications (Fig. 4**c**), **Eubiota** consistently nominated LacCom as the superior consortium design. The rationale emphasized SCFA production and *Lactobacillaceae*-associated functions supporting epithelial barrier protection and anti-inflammatory signaling [102, 103]. In contrast, AceCom was deprioritized due to inclusion of *R. gnavus*, a taxon associated with pro-inflammatory dysbiosis [104, 105]. Whereas frontier general-purpose LLMs [106–108] exhibited inconsistent consortium selection, **Eubiota** maintained stable, evidence-grounded prioritization (Extended Data Fig. 6).

#### *In vivo* validation of Eubiota-guided consortium selection

We tested these predictions in a DSS-induced colitis mouse model (2.5% w/v) following the **Eubiota**-generated protocol (Fig. 4**d**, Supplementary Note 6). Both AceCom and LacCom attenuated DSS-induced weight loss (Fig. 4**e**); however, LacCom-treated group transitioned more rapidly into recovery, resulting in significantly lower Disease Activity Index (DAI) scores by day 10 (Fig. 4**f**). Consistent with these findings, LacCom substantially rescued inflammation-associated colon shortening, whereas AceCom did not (Fig. 4**g**). Longitudinal fecal lipocalin-2 (LCN-2), a sensitive biomarker of intestinal inflammation, was significantly reduced by day 10 in LacCom-treated mice, while remaining elevated in AceCom-treated animals (Fig. 4**h**). These results indicate that LacCom not only mitigates acute inflammatory injury but also accelerates resolution in this model of colitis.

#### *De novo* consortium design under laboratory constraints

To assess generalization beyond predefined candidates, we restricted **Eubiota** to a laboratory inventory of 22 strains (Supplementary Note 6) and tasked it with *de novo* consortium design. The system proposed a four-member community (*Blautia hansenii, Bifidobacterium breve, Faecalibacterium prausnitzii*, and *Roseburia intestinalis*) integrating complementary functions in epithelial barrier support, acetate and lactate production, and butyrate-mediated immunomodulation [109–112] (Fig. 4**i**). Following the **Eubiota**-generated protocol, we evaluated this synthetic community (SynCom) in the DSS colitis model (Fig. 4**j**). By day 10, SynCom significantly attenuated weight loss and DAI scores (Fig. 4**k, l**), partially rescued colon shortening (*P* = 0.054) (Fig. 4**m**), and reduced fecal LCN-2 levels relative to DSS-only controls (Fig. 4**n**). Altogether, these findings demonstrate that **Eubiota** can translate ecological principles into experimentally testable therapeutic consortia that mitigate intestinal inflammation.

### 3.4 Eubiota-enabled pathogen-biased antibiotic cocktail design

#### Rational assembly of antibiotic cocktails

Effective antimicrobial therapy in the gut requires suppressing pathogens and pathobionts while preserving beneficial commensals, a balance that remains difficult to achieve with existing antibiotics. Although recent LLM-based approaches have advanced *de novo* antimicrobial design [113, 114], translation remains limited by synthesis cost and regulatory burden [115]. We therefore tasked **Eubiota** with designing a selective cocktail to inhibit three common pathobionts, *Pseudomonas aeruginosa* (*Pa*), *Citrobacter portucalensis* (*Cp*), and *Klebsiella pneumoniae* (*Kp*) while sparing representative commensals including *E. coli* (*Ec*), *Bacteroides* species, and *Bifidobacterium* species. **Eubiota** executed a three-stage workflow (Fig. 5**a**) that automated antibiotic cocktail selection, extraction of MIC data to define dosing ranges, and (Phase III) generation of executable protocols (Supplementary Note 6).

**Figure 5:**
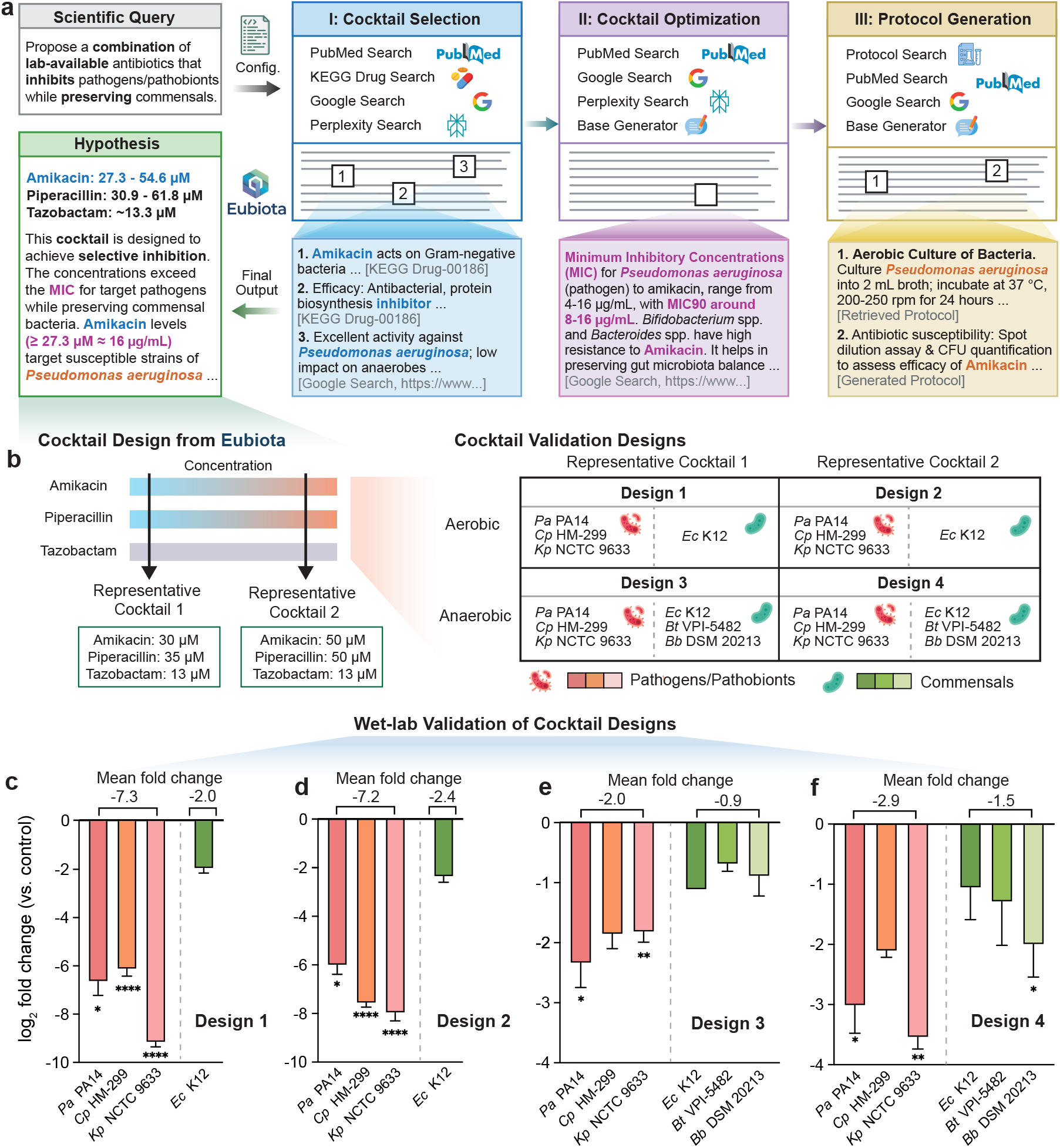
Rational design of pathogen-biased antibiotic cocktails. **a**, Agentic workflow for selective antibiotic cocktail design. **Eubiota** integrates pathogen susceptibility profiles, resistance mechanisms, and drug-target pathway annotations using literature and database tools (PubMed, KEGG, MDIPID) alongside general web search. The modular pipeline comprises three stages: (I) Cocktail Selection, (II) MIC-guided Concentration Optimization, and (III) Protocol Generation. The resulting hypothesis proposes a combination of amikacin, piperacillin, and tazobactam with concentrations selected to exceed pathogen MICs while minimizing inhibitory effects on representative commensals. **b**, Two representative cocktails spanning the lower (**Cocktail 1**) and upper (**Cocktail 2**) bounds of the **Eubiota**-predicted selective concentration window were evaluated. Cocktails were tested against *P. aeruginosa* PA14 (*Pa* PA14), *K. pneumoniae* NCTC 9633 (*Kp* NCTC 9633), *C. portucalensis* HM-299 (*Cp* HM-299), and *E. coli* K12 (*Ec* K12) under aerobic and anaerobic conditions. Obligate anaerobes *B. thetaiotaomicron* VPI-5482 (*Bt* VPI-5482) and *B. breve* DSM 20213 (*Bb* DSM 20213) were assessed under anaerobic conditions only. **c–f**, Experimental validation of antibiotic efficacy and selectivity. **c, d**, Aerobically grown stationary-phase cultures (37 °C, 24 h) were diluted 1:2 and exposed to antibiotic cocktails for 24 h. Viability is shown as log_2_ fold change in CFU/mL relative to vehicle controls. Preservation of *Ec* K12 viability indicates selective sparing under aerobic conditions. **e, f**, Under anaerobic conditions, pathogen suppression was reduced, yet commensal viability was partially preserved. Numbers above bars indicate the mean log_2_ fold change for pathogen (red) and commensal (green) groups. Data are mean ± s.e.m. (*n* = 3 biological replicates). Statistical significance was determined by two-tailed Student’s *t*-test on raw CFU values comparing antibiotic-treated and untreated controls. ^∗^*P* < 0.05, ^∗∗^*P* < 0.01, ^∗∗∗^*P* < 0.001, ^∗∗∗∗^*P* < 0.0001.

#### Inventory-constrained design reveals baseline selectivity

Under laboratory inventory limits, **Eubiota** identified a design including piperacillin, tazobactam, amikacin and a concentration window (amikacin 27.3-54.6 *µ*M, piperacillin 30.9-61.8 *µ*M, tazobactam ∼13.3 *µ*M) via multi-agent synthesis of literature and database evidence [116–118]. To assess **Eubiota**’s recommendations, we selected two formulations in the high and low range and tested their effects using a high-density outgrowth assay (Fig. 5**b**). Stationary-phase cultures were diluted 1:2 into fresh medium pre-supplemented with antibiotics. Under aerobic conditions, both designs achieved robust suppression of the target pathogens, while the commensal *Ec* K12 remained relatively tolerant (Fig. 5**c, d**). We subsequently evaluated the cocktails under anaerobic conditions to simulate the distal gut environment. Pathogen suppression was reduced, with partial sparing of *Ec* K12, *Bacteroides thetaiotaomicron* (*Bt*) VPI-5482, and *Bifidobacterium breve* (*Bb*) DSM 20213 (Fig. 5**e, f**). This attenuation is consistent with the reduced bioenergetic uptake of aminoglycosides and diminished growth-dependent activity of *β*-lactams under anaerobiosis [119, 120]. These results exhibit the system’s logical rigor while highlighting the inherent complexity of biological questions surrounding antibiotic treatment and gut microbiota [121, 122].

One possible way for further optimization is the integration of iterative scientist feedback. In an unconstrained search, **Eubiota** proposed a four-drug cocktail (piperacillin, tazobactam, amikacin, and ciprofloxacin) (Extended Data Fig. 7**a**). The initial cocktail achieved stronger pathogen clearance meanwhile substantial commensal loss when administrated at bacteria inoculation (Extended Data Fig. 7**b, d**). We then implemented a workflow using this wet-lab outcome and expert feedback: initial concentrations are toxic to commensals. In response, **Eubiota** recalibrated the dosage (Extended Data Fig. 7**c**). This refinement process improved the final outcome, as it preserved some commensal populations at viable levels while maintaining pathogen inhibition (Extended Data Fig. 7**d, e**). This sparing may be attributed to the commensals entering a tolerant, non-replicative state and potential pharmacological antagonism between the cocktail components [123–126]. Despite the challenges of this biological setup, these findings demonstrate the system’s capacity to reason through complex scientific questions and iteratively optimize proposed strategies.

### 3.5 Eubiota guided anti-inflammatory molecule discovery

#### Prioritizing immunomodulatory metabolites from human dietary interventions

Human dietary intervention studies provide a powerful but challenging substrate for immunomodulatory discovery [127–129]. High inter-individual variability and heterogeneous metabolite annotation often obscure causal links between diet and immune phenotypes (Fig. 6**a**). To navigate this high-dimensional search space, we leveraged metabolomic data from the Twins Nutrition Study (TWIN; NCT05297825), a controlled trial comparing vegan and omnivorous diets in identical twins [130, 131]. We tasked **Eubiota** with screening 397 serum metabolites enriched in the vegan arm for potential NF-*κ*B inhibitory activity. **Eubiota** integrated literature and database retrieval (Extended Data Fig. 4**c, d**) with mechanistic and phenotypic reasoning to assign each metabolite a structured likelihood score. These scores were aggregated across runs to generate a ranked consensus, and the top candidates were translated into executable NF-*κ*B reporter assay protocols (Fig. 6**b**).

**Figure 6:**
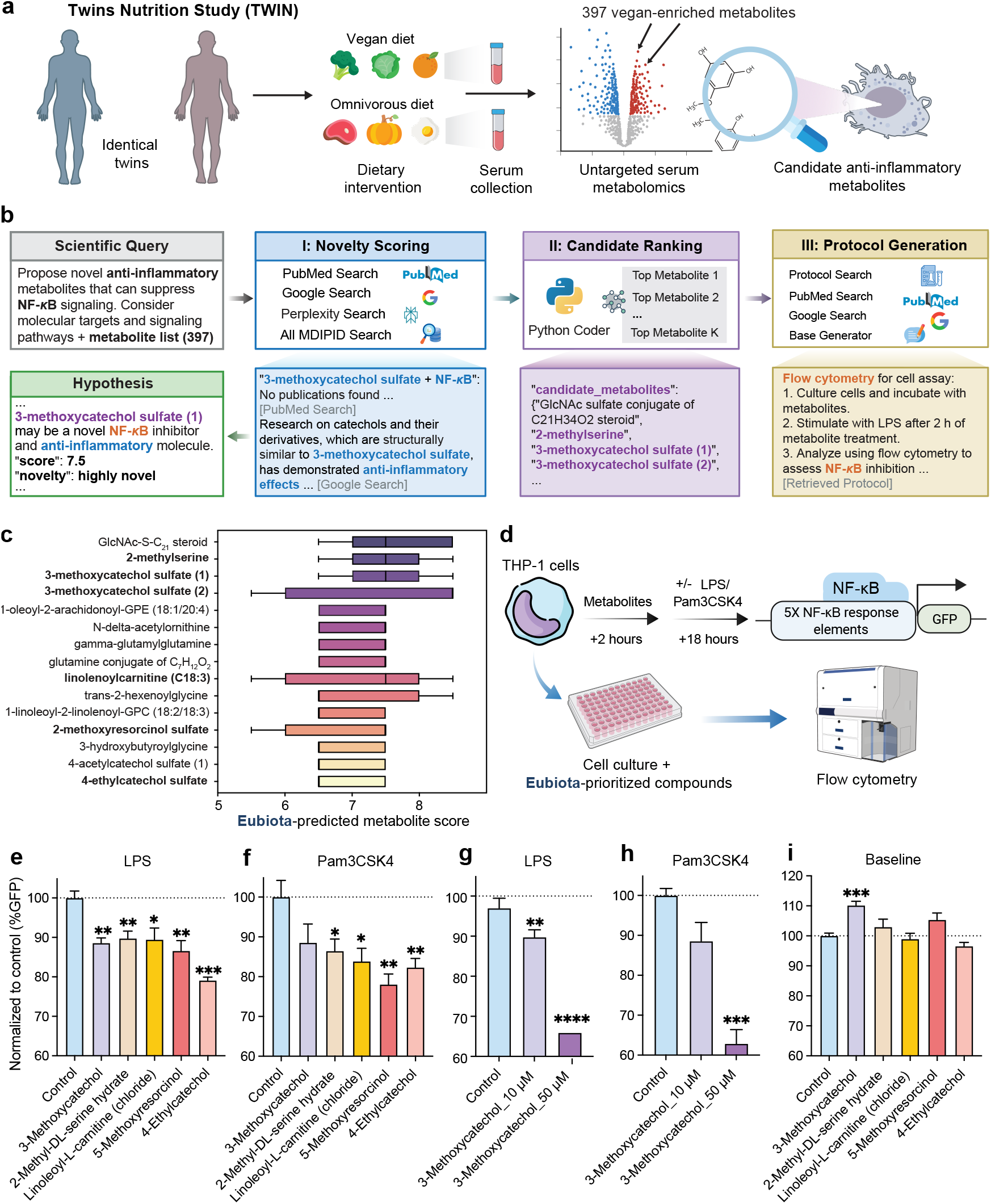
Eubiota-driven identification of novel immunomodulatory metabolites from human cohorts. **a**, Study design leveraging untargeted metabolomics data from the Twins Nutrition Study (TWIN) [131]. Identical twin pairs were assigned to different diets (vegan versus omnivorous). The metabolomics data has the potential for identification of circulating metabolites enriched under the vegan dietary intervention associated with anti-inflammatory activity. **b, Eubiota** discovery workflow for metabolite prioritization. A set of 397 vegan-enriched circulating metabolites from the TWIN serum metabolomics dataset [131] was evaluated using the full tool suite. Metabolites were ranked based on composite scores integrating predicted anti-inflammatory potential and bibliometric novelty. The system also generated a literature-grounded experimental protocol for *in vitro* validation of top-ranked candidates using cell-based assays. **c**, Top-ranked metabolites based on **Eubiota**-assigned composite scores aggregated across five independent runs. **d**, *In vitro* validation schematic. THP-1 NF-*κ*B reporter cells were pre-treated with candidate metabolites, stimulated with lipopolysaccharide (LPS) or Pam3CysSerLys4 (Pam3CSK4, a lipopeptide), and NF-*κ*B activity was quantified by GFP fluorescence via flow cytometry after 18 h. **e–i**, Experimental validation of prioritized metabolites in THP-1 NF-*κ*B reporter cells. **e, f**, Suppression of inflammatory signaling. All five prioritized candidates (10 *µ*M) significantly reduced NF-*κ*B activation induced by **e**, LPS and **f**, Pam3CSK4. **g, h**, Dose-response analysis of 3-methoxycatechol (10 *µ*M and 50 *µ*M) demonstrating concentrationdependent reduction of NF-*κ*B signaling under **g**, LPS and **h**, Pam3CSK4 stimulation. **i**, Baseline NF-*κ*B activity in the absence of inflammatory stimulation. 3-methoxycatechol increased baseline GFP expression relative to solvent control. All values are normalized to solvent controls (dotted line) and shown as mean ± s.e.m. (*n* = 4 biological replicates). Statistical significance was determined by two-tailed Student’s *t*-test comparing median fluorescence intensity of compound-treated samples to solvent controls. ^∗^*P* < 0.05, ^∗∗^*P* < 0.01, ^∗∗∗^*P* < 0.001, ^∗∗∗∗^*P* < 0.0001.

#### Prioritized metabolites reveal convergent biochemical themes

The top-ranked metabolites clustered into recurring biochemical classes (Fig. 6**c**), including sulfated phenolics (3-methoxycatechol sulfate, 4-ethylcatechol sulfate, 4-acetylcatechol sulfate, 2-methoxyresorcinol sulfate), nitrogen-containing amino acid derivatives (2-methylserine, N-*δ*-acetylornithine), and lipid-associated conjugates (linolenoylcarnitine [C18:3], linoleoylcarnitine [C18:2]). The highest-scoring feature was a GlcNAc-sulfate conjugate of a C_21_H_34_O_2_ steroid. Despite chemical diversity, aromatic sulfate conjugates and modified lipid species emerged as recurrent structural motifs within the veganenriched metabolome, suggesting convergent biochemical signatures associated with anti-inflammatory potential.

#### Experimental validation confirms anti-inflammatory activity

We selected five commercially available parent or analog compounds for validation: 3-methoxycatechol, 2-methyl-DL-serine hydrate, linoleoyl-L-carnitine, 5-methoxyresorcinol, and 4-ethylcatechol. Their effects were assessed following TLR4 (lipopolysaccharide [LPS]) or TLR1/2 (Pam3CysSerLys4 [Pam3CSK4]) stimulation, using a human THP-1 monocytic cell line engineered to express Green Fluorescent Protein (GFP) under the control of NF-*κ*B, a key transcriptional regulator of the innate immune response [132, 133] (Fig. 6**d**). The experiments were guided by **Eubiota** proposed protocols (Supplementary Note 6). All five metabolites significantly reduced reporter expression at physiologically relevant concentrations (10 *µ*M to 500 *µ*M, *P* < 0.05) [134–137] (Fig. 6**e, f**). 3-methoxycatechol exhibited a dose-dependent suppression of NF-*κ*B signaling (Fig. 6**g, h**), although it induced modest baseline expression in the absence of stimulation (Fig. 6**i**). This dual phenotype is consistent with catechol auto-oxidation and reactive oxygen species generation [138, 139], a behavior characteristic of redox-active natural products [140]. While 4-ethylcatechol has been reported to suppress NF-*κ*B in murine cells [134], direct evidence for several of the remaining compounds was previously lacking. Notably, although 3-methoxycatechol sulfate has been associated with interleukin-1 beta modulation [141], our findings provide, to our knowledge, the first demonstration that the parent compound itself exhibits the suppression of NF-*κ*B reporter signaling. Importantly, prior studies have linked 3-methoxycatechol to activation of GPR35, a G protein–coupled receptor with anti-inflammatory roles in macrophages and the intestinal immune system [142, 143]. It suggests that this metabolite may impact immune cells through multiple, partially independent signaling pathways beyond NF-*κ*B regulation. Collectively, these results expand the repertoire of diet-associated host–microbe co-metabolites with demonstrable anti-inflammatory activity and illustrate how agentic prioritization can convert heterogeneous human metabolomic data into experimentally validated mechanistic insights.

#### Reliability, stability, and tool-grounded discovery

To assess robustness, we executed the full workflow across five independent runs. Score distributions were consistent, with moderately strong concordance (mean Pearson *r* = 0.57) (Supplementary Fig. 4**c**). ICC analysis indicated moderate reliability for single runs (ICC(2,1) = 0.57) and high reliability when aggregating across runs (ICC(2,5) = 0.87), exceeding permutation-based null expectations (Supplementary Fig. 4**d, e**). Tool usage patterns and citation sources aligned with task-specific requirements (Extended Data Fig. 4**c, d**), indicating that **Eubiota** predictions were grounded in structured evidence rather than uncontrolled generative variability. Together, these results demonstrate that **Eubiota** reliably identifies biologically meaningful and experimentally actionable immunomodulatory signals from heterogeneous human intervention metabolomics datasets.

### 3.6 Human evaluation study

To evaluate real-world utility, we conducted a randomized, double-blinded comparison between **Eubiota** and a frontier general-purpose model (GPT-5.1 [50]). Fifteen domain experts in microbiology and immunology assessed anonymized, shuffled responses to open-ended queries ranging from general microbiome concepts to specific mechanistic research questions (representative examples are provided in Supplementary Note 5). Across 163 independent evaluations, **Eubiota** was preferred in 70.6% of cases (*n* = 115), compared to 25.8% for GPT-5.1 (*n* = 42), with 3.7% ties (*n* = 6) (Fig. 1**d**).

Qualitative feedback revealed that **Eubiota**’s advantage was primarily attributable to its evidence-grounding reasoning and transparency. Participants emphasized the importance of comprehensive source integration and verifiable citations for academic trust and experimental planning. GPT-5.1 was occasionally favored for its concise synthesis and linguistic fluency; however, **Eubiota** was more consistently preferred for research-intensive queries requiring methodological rigor and verifiable support. Together, these results suggest that modular agentic systems offer practical advantages for scientific workflows in which auditability and evidence-based traceability are essential.

## 4 Discussion

**Eubiota** introduces a structured paradigm for integrating agentic AI into biological discovery by decoupling planning, tool execution, verification, and grounded synthesis into specialized modules coordinated through shared memory. This architecture transforms scientific reasoning from “black box” generation into an auditable, state-aware process in which intermediate goals, evidence, and decisions are explicitly recorded. This design supports step-by-step refinement of objectives and constraints, while preserving continuity throughout multi-turn exploration. The resulting modularity supports generalization to microbiome-focused mechanistic tasks while enabling extensibility to the resources and laboratory protocols of new domains. Crucially, outcome-driven reinforcement learning enhances reliability. Optimizing the Planner policy with outcome-based rewards (using GRPO-MAS) promotes evidence-seeking and cross-checking behaviors that reduce compounding errors in multi-turn workflows. Rather than optimizing isolated responses, this approach aligns long-horizon reasoning with verifiable outcomes, strengthening logical coherence under domain constraints.

Beyond methodological contributions, **Eubiota** discovered experimentally validated biological insights into host-microbiome interactions. It identified the *uvr–ruv* DNA repair axis as a determinant of bacterial fitness under nitrosative stress, highlighting that genome maintenance is essential for persistence in inflammatory environments. The enrichment of these genes in human IBD metagenomes suggests that strain-level adaptation to host-associated stress can potentially be observed on the molecular scale, which has implications for diagnostics and therapeutic intervention. At the community scale, **Eubiota**’s *de novo* design of a four-member consortium demonstrates a path toward mechanistically defined, probiotic-like alternatives to FMT. Pathogen-biased antibiotic optimization further illustrates how iterative human feedback can refine complex biological outputs under realistic constraints. Finally, the system’s prioritization of vegan diet-enriched metabolites uncovered multiple suppressors of NF-*κ*B activity, providing mechanistic support for diet-associated immunomodulation and expanding the catalog of host-microbe cometabolites with anti-inflammatory potential. Collectively, these applications demonstrate that modular agentic workflows can move beyond descriptive aggregation of existing knowledge to generate experimentally actionable new hypotheses across molecular and community scales.

The modular design of **Eubiota** offers a scalable foundation for broader scientific deployment. Future iterations may improve efficiency through lightweight verifier policies, adaptive early stopping, caching of retrieved evidence, and uncertainty-triggered selective reruns to reduce computational overhead while maintaining auditability. Reliability could be further enhanced through prioritization of up-to-date sources for time-sensitive evidence, automated cross-checking when data are sparse or conflicting, and explicit rewards for reproducible reasoning traces under fixed compute budgets.

Scientifically, the **Eubiota** framework enables a transition toward integrated, multi-modal discovery. Future work may link enrichment and fitness-associated signals to context-specific causality by combining data from strain-resolved transcriptomics, targeted perturbations, and host-level validations. Therapeutic design can be further refined under explicit environmental and host constraints, including diet, baseline microbiota composition, host genetics, and inflammatory state, to distinguish generalizable design principles from context-dependent requirements. To elucidate the mechanisms of antibiotics and anti-inflammatory metabolites, future iterations can incorporate comprehensive *in vitro* assays and *in vivo* models. Ultimately, **Eubiota** positions modular agentic systems not as replacements for scientific expertise, but as interactive collaborators that scale evidence integration and accelerate the translation of complex biological data into discovery and mechanistic understanding.

## Extended Methods

### Toolset implementation and details

#### Tool construction framework

All **Eubiota** tools inherit from a unified `Tool` base class that standardizes interfaces across heterogeneous capabilities (Extended Data Fig. 1**a**). Each tool specification defines structured metadata, including: (1) `name` and `description` defining the tool’s purpose and supporting Planner selection; (2) input parameters (`input_kwargs`) specifying required and optional arguments with explicit type constraints; (3) an `output_schema` specifying the structure of returned data; (4) documented `limitations` describing known constraints and failure modes; and (5) `best_practices` guidelines for reliable usage.

The Executor dynamically instantiates tools according to the Planner’s selection and translates planned actions (including required context and expected outputs) into executable tool-call commands. Inputs are formatted and checked against declared schemas and argument constraints prior to execution. To ensure robustness, the Executor implements a schema-aware iterative correction loop: when validation errors occur, structured error feedback is used to repair and regenerate arguments in subsequent attempts (Extended Data Fig. 1**b**).

#### Context retrieval and synthesis

Tools such as URL Context Search and MDIPID Search retrieve passages from web pages or database entries. Unlike retrieval-only frameworks that return raw context [35], these tools generate concise, evidence-grounded summaries to increase information density and reduce latency. To enable this, we implemented a retrieval-augmented generation (RAG) pipeline (Extended Data Fig. 1**c**). Source corpora are segmented into ∼200-word fixed-length chunks, and both the input query and text segments are embedded into 3,072-dimensional semantic vectors using the OpenAI text-embedding-3-large model^1^. The top-*k* chunks (default *k* = 10) are retrieved based on cosine similarity, after which an LLM (e.g., GPT-4o [60]) synthesizes a compact response grounded in the retrieved evidence. This design preserves explicit traceability to source material while reducing prompt length, thereby improving the robustness and efficiency of tool-mediated scientific reasoning.

#### Web search tools

To support up-to-date information retrieval, we integrated a **Google Search** tool using the Grounding with Google Search feature^2^. This integration provides access to real-time web content, thereby mitigating hallucinations, and enables citation-based responses for information beyond the model’s training cutoff. In parallel, **Perplexity Search**^3^ retrieves ranked web results with aggregated citations and summaries. For encyclopedic content, Wikipedia Search queries the Wikipedia API^4^ and uses RAG to summarize entries. Finally, to allow fine-grained inspection of specific sources identified during searches, the **URL Context Search** tool extracts full-text content and synthesizes relevant passages using RAG.

#### Literature search tool

We implemented a **PubMed Search** tool that interfaces with the NCBI Entrez E-utilities API^5^ to retrieve article metadata (titles, abstracts, and PMIDs) in response to keyword queries. Retrieved records are processed by an LLM (e.g., GPT-4o [60]) to generate concise, citation-linked summaries with in-text references (formatted as “[1] PMID:XXXXXX”). This design ensures explicit traceability from generated statements to primary literature sources.

#### KEGG database tools

Kyoto Encyclopedia of Genes and Genomes (KEGG) tools query the GENES, GENOME, Drug, and Disease databases^6^. The **KEGG Gene** and **Organism Search** tools access the GENES and GENOME databases to retrieve structured annotations, including gene functions, metabolic pathways, and taxonomic information. The **KEGG Drug** and **Disease Search** tools retrieve chemical structures, drug-target relationships, and disease-associated pathway mappings. All tools return structured JSON outputs, enabling the Planner to systematically map mechanistic relationships between microbial genes, pharmacological interventions, and host-relevant pathways.

#### MDIPID database tools

To support structured querying of complex microbe–drug–disease relationships, we integrated the Microbiota-Drug Interaction and Disease Phenotype Interrelation Database (MDIPID)^7^. In contrast to the structured queries used for KEGG, the **MDIPID Gene, Microbe**, and **Disease Search** tools use retrieval-augmented generation (RAG) over pre-indexed database profiles. Queries are processed using a hybrid retrieval strategy that combines exact string matching with vector similarity search, after which an LLM synthesizes responses grounded in curated functional annotations and interaction evidence.

#### Specialized analysis tools

We equipped **Eubiota** with specialized modules linking computational reasoning, experimental data, and laboratory practice. The **Gene Phenotype Search** tool retrieves pre-computed cumulative fitness scores derived from 118 human gut bacterial growth screens under nitric oxide (NO). Growth fitness was quantified as the differential area under the curve (ΔAUC) relative to untreated controls using the Baranyi-Roberts model [70, 71]. Gene-level cumulative scores were then computed across characterized orthologous gene groups to identify conserved determinants of nitrosative stress for integration into the **Eubiota** discovery workflow. This tool enables rapid, data-driven candidate prioritization.

To support experimental design, the **Protocol Search** tool retrieves detailed procedures from a curated library of 55 assays spanning molecular biology, microbiology, and metabolomics. In addition, the **Doc Context** and **Database Context Search** tools implement RAG over user-supplied documents and structured files, thereby enabling reasoning over custom datasets without retraining.

#### Generation and computation tools

Two utility tools support numerical computation and internal reasoning. The **Python Coder** generates and executes code within a sandboxed environment, enabling statistical analysis, data processing, and precise numerical computation beyond the native arithmetic capabilities of language models. The **Base Generator** tool uses the underlying LLM for synthesis, planning, and content creation in the absence of external retrieval.

### Planner optimization

#### Reinforcement learning framework

We parameterize the **Eubiota** Planner as a learnable policy and optimize it using reinforcement learning (RL) within the full multi-agent execution loop. Planning decisions, such as tool selection and subgoal specification, are made sequentially across multiple turns, and early actions can influence downstream outcomes. Unlike supervised fine-tuning, which optimizes individual steps independently, our approach trains the Planner on-policy with respect to trajectory-level outcomes. Building on findings that RL can improve reasoning by encouraging self-correction and long-horizon planning [37, 39, 144], we develop the Group Relative Policy Optimization for Multi-Agent Systems (GRPO-MAS) strategy, a variant of GRPO [57] adapted for multi-agent reasoning (Extended Data Fig. 2**a**). GRPO-MAS addresses sparse, delayed rewards by assigning a trajectory-level outcome signal and propagating credit through the sequence of planning decisions, enabling optimization of long-horizon tool-use behavior.

#### Markov decision process and reward signal

We model the discovery workflow as a Markov decision process in which the Planner is parameterized as a stochastic policy *π*_*θ*_ (*a*_*t*_ | *S*_*t*_). At each turn *t*, the policy selects an action *a*_*t*_ (comprising subgoal specification and tool selection) conditioned on the current state *S*_*t*_, defined as the fixed query *q* together with the accumulated shared memory. For a given query *q*, the system samples a group of *G* independent execution trajectories (rollouts) {*τ*_1_, …, *τ*_*G*_} from the current policy 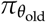. Each trajectory *τ*_*i*_ proceeds for *T*_*i*_ turns and terminates either when the Verifier signals sufficiency or when the maximum turn limit is reached. Upon termination, the Generator produces a final evidence-grounded response. A terminal binary reward *R*(*τ*_*i*_) ∈ 0, 1 is assigned to each trajectory based on agreement between the generated answer and the ground-truth label.

#### Group-relative advantage estimation

To stabilize training and reduce variance without introducing a separate value network, we compute a group-relative advantage *A*_*i*_ for each trajectory *τ*_*i*_. This normalizes the reward of an individual rollout relative to the group baseline:

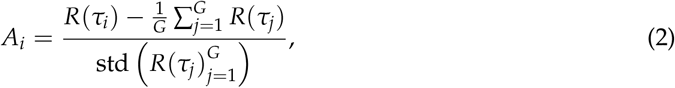

where std 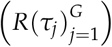 denotes the standard deviation of rewards within the rollout group. Crucially, the trajectory-level advantage *A*_*i*_ is assigned uniformly to all Planner-generated tokens within *τ*_*i*_, providing a shared credit signal for the sequence of planning decisions and tool selections that produced the final outcome. Positive advantages increase the likelihood of actions associated with successful trajectories, whereas negative advantages suppress actions associated with lower-performing rollouts.

#### Optimization objective

We optimize the policy parameters *θ* by maximizing a clipped surrogate objective with Kullback–Leibler (KL) regularization that encourages high-advantage actions while constraining policy divergence. Specifically, we employ a token-level Proximal Policy Optimization (PPO) objective averaged across the group [145]:

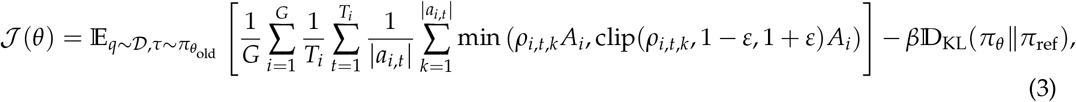

where |*a*_*it*_| denotes the token length of action *a*_*it*_, and *ρ*_*it*_ is the probability ratio 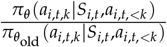 for the *k*-th token of action *a*_*i,t*_. The clipping function clip (·, 1 − *ε*, 1 + *ε*) restricts the magnitude of policy updates, and the KL divergence term D_KL_ (weighted by *β*) regularizes the learned policy toward a reference policy *π*_ref_ to control drift. This objective converts sparse, trajectory-level outcomes into token-level learning signals that enable credit assignment to intermediate planning decisions throughout the multi-turn workflow.

#### Planner optimization and training parameters

We optimized the **Eubiota Planner** using Group Relative Policy Optimization for Multi-Agent Systems (GRPO-MAS), while keeping the Executor, Verifier, and Generator fixed. Training employed a learning rate of 1 × 10^−6^, a batch size of 32, and *G* = 8 rollouts per query. Planner actions were sampled with a temperature 0.7 to promote exploration of tool selections and multi-turn strategies. Policy updates were regularized with a KL-divergence penalty relative to a frozen reference policy (*β* = 0.001). The Planner’s maximum generation length was set to 2,048 tokens, and each rollout was limited to three Planner– Executor–Verifier cycles to balance exploration depth with training throughput. Upon termination, the Generator produced an evidence-grounded response, which received a terminal reward from an LLM-based evaluator. The evaluator assessed agreement between the generated answer and the ground-truth label for each query; GPT-4o [60] was used for reward evaluation due to its high throughput and low latency. To reduce nondeterminism and isolate planning variability, tool-invoking LLMs operated at temperature 0.0 during training. Optimization was performed on four NVIDIA A100 GPUs.

#### Training dataset construction

To equip **Eubiota** with microbiome domain expertise while retaining general reasoning and retrieval capabilities, we constructed a heterogeneous training corpus of 2,000 labeled instances. The dataset was composed of four components mixed at a 6:2:1:1 ratio across microbiome reasoning, retrieval, general biomedical knowledge, and mathematical reasoning (Extended Data Fig. 2**b**). Microbiome-specific examples were derived from the Microbiota-Drug Interaction and Phenotypic Insights Database (MDIPID) and constituted the core of the dataset. Selected subsets covering disease phenotypes and microbiota–disease associations were reformulated as multiple-choice inference tasks requiring mechanistic reasoning. To broaden generalization, we incorporated three auxiliary sources: (i) fact-based retrieval tasks from Natural Questions (NQ) [146] to train query formulation and retrieval triggering; (ii) biomedical question-answering examples from PubMedQA [147] and MedQA [148] to provide domain-general context; and (iii) mathematical reasoning problems from DeepMath-103K [149] to encourage multi-step inference. Detailed preprocessing, quality-control (including deduplication and leakage prevention), and task templates are provided in Extended Data Fig. 2**b** and Supplementary Note 5.

### Inflammatory stress-responding gene discovery

#### Genetic discovery workflow

To identify gene-phenotype associations underlying nitrosative stress tolerance, we employed **Eubiota** in a four-phase discovery workflow to screen bacterial genes for potential fitness advantages under NO exposure.

- Phase I (Likelihood Scoring): Each candidate gene was evaluated independently using a composite scoring rubric integrating scientific relevance, reported phenotypic evidence, known or inferred molecular function, and mechanistic plausibility in oxidative or nitrosative stress responses. Evidence was drawn from quantitative phenotypic data and literature-derived annotations. Broadly conserved housekeeping genes were deprioritized to enrich for stress-adaptive or conditionally advantageous functions.
- Phase II (Gene Ranking): Genes were ranked by composite score, and the top 10 candidates were selected for downstream validation. Data parsing and rank-sorting were automated using the Python Coder tool.
- Phase III (Protocol Generation): For each prioritized gene, **Eubiota** generated gene-specific experimental protocols by assembling standardized microbiological and genetic assays via the Protocol Search tool. Targeted PubMed queries were executed to incorporate gene-specific contextual information, enabling parameter refinement tailored to the individual genetic background.
- Phase IV (Transposon Mutant Mapping): Prioritized genes were mapped to corresponding mutants within an arrayed *B. breve* transposon library to facilitate validation. Mapping and metadata parsing were automated using the Python Coder tool to ensure accurate alignment between *in silico* predictions and available mutant stocks.

In each workflow execution, every gene was assigned a likelihood score (1–100) together with a structured rationale grounded in tool-derived evidence. The full workflow was repeated five times independently to ensure reliable gene scores, and final rankings were determined by the mean score across runs.

#### Stability and tool transition analysis

To assess reproducibility of gene scoring and characterize tool utilization patterns, we performed two complementary analyses. First, we examined the distribution of gene likelihood scores across five independent workflow runs using ridgeline density plots generated in R with `ggplot2` and `ggridges`. Kernel density estimates were overlaid for each run [150] to enable direct comparison of score distributions and evaluate cross-run consistency in distributional shape, variance, and range. Second, we analyzed tool usage dynamics by constructing directed transition graphs from parsed execution logs. Nodes correspond to individual tools, and directed edges represent sequential transitions between tool invocations within a workflow. Edge weights reflect cumulative transition frequencies aggregated across runs. These graphs quantify recurrent tool sequences, stochastic branching patterns, and the task-specific logic inherent in **Eubiota**’s decision-making process.

#### Intraclass correlation coefficients (ICCs) analysis

ICCs were computed to assess the reliability of gene likelihood scores across independent workflow runs [151]. We applied a two-way randomeffects model with absolute agreement, treating both genes and runs as random effects [152]. This model is appropriate because each run constitutes a stochastic realization of the same agentic pipeline, and the objective was to evaluate consistency of gene-level scores across runs. We report both the single-measure ICC (ICC(2,1)), which quantifies the reliability of an individual run, and the average-measure ICC (ICC(2,*k*)), which estimates the reliability of the mean score aggregated across *k* runs. The coefficients are defined as:

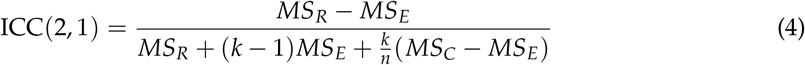

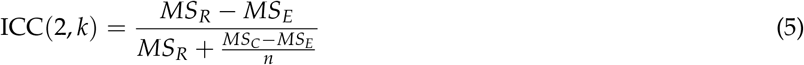

where *MS*_*R*_ denotes the mean square for rows (between-gene variance), *MS*_*C*_ the mean square for columns (between-run variance), *MS*_*E*_ the residual mean square, *k* the number of runs, and *n* the number of genes.

#### Candidate gene selection and functional mapping

To quantify phenotypic responses across the human gut microbiota, 118 representative strains from the hCom1 and hCom2 communities [69] were cultured anaerobically (85% N_2_, 10% CO_2_, 5% H_2_) for 48–72 h. Pre-cultures were subcultured (1:100) into 384-well plates with or without 10 *µ*M DETA NONOate, and growth kinetics were monitored at 37 °C using an Epoch2 microplate reader (BioTek) with double orbital shaking. Growth curves were fitted to the Baranyi–Roberts model [70, 71], and fitness was quantified as the differential area under the curve (ΔAUC) relative to untreated controls. To enable high-confidence functional mapping, analysis was restricted to protein-coding sequences annotated with standardized gene symbols by the NCBI Prokaryotic Genome Annotation Pipeline. Orthologous genes were collapsed across strains into shared functional units, while uncharacterized, locus-tag-specific hypothetical proteins were excluded. For each ortholog, a cumulative fitness score was computed as the sum of normalized ΔAUC values across all strains harboring that gene, thereby identifying conserved genetic determinants of nitrosative stress tolerance. The ranked ortholog dataset was incorporated into the Gene Phenotype Search tool and used as input for the **Eubiota** discovery workflow.

#### Gene-phenotype experimental validation

To validate gene-phenotype associations, we used an arrayed, barcoded transposon mutant library of *B. breve* UCC2003 [76]. Mutant strains were cross-referenced to the reference genome using an indexed insertion map. For each prioritized gene, **Eubiota** mapped candidate loci to corresponding mutant identifiers, enabling retrieval of loss-of-function strains for validation. Prior to phenotyping, mutants and wild-type (WT) controls were revived from glycerol stocks and cultured anaerobically at 37 °C for 48 h in pre-reduced, cysteine-free Yeast Casitone Fatty Acid with Carbohydrate (YCFAC) medium. Starter cultures were diluted 1:100 into fresh medium with or without 10 *µ*M DETA NONOate (Cayman Chemical), used as an NO donor. Because NO is highly reactive and short-lived, 10 *µ*M DETA NONOate was used to approximate sustained exposure to 10 *µ*M NO. Growth kinetics were measured in 96-well clear, flat-bottom plates sealed with transparent lids and incubated at 37 °C with double orbital shaking in an Epoch2 microplate reader (BioTek). Raw optical density at 600 nm (OD_600_) values were background-corrected by subtracting the initial reading of each well and analyzed using a published pipeline [77] to calculate area under the curve (AUC).

#### Validation of gene enrichment in public IBD datasets

To evaluate the clinical relevance of **Eubiota**-prioritized genes, we performed cross-cohort enrichment analyses using two independent human metagenomic resources. First, we analyzed a curated cross-study meta-analysis of fecal metagenomes from patients with inflammatory bowel disease (IBD) and healthy controls [78]. Prioritized genes were mapped using KEGG Orthology (KO) identifiers, and relative enrichment was quantified using generalized fold change (gFC) values reported across cohorts in the study.

Second, we analyzed a strain-resolved genomic resource comprising genotype reconstructions from approximately 140,000 gut microbial strains [79]. We performed an intersection analysis to identify genes that (i) were reported in the strain-level dataset [79] and (ii) ranked within the top 50 genes generated by the **Eubiota** pipeline. Differential enrichment between IBD-adapted and health-associated strains was assessed using false discovery rate (FDR)-adjusted *P* values as reported in the source publication. Full dataset details and statistical methodologies are described in the respective original studies [78, 79].

### Therapeutic design for colitis treatment

#### Therapy design workflow

The **Eubiota** framework was evaluated across three microbial community design tasks. As an initial validation, the system was challenged with a multiple-choice selection task to identify a canonical butyrate-producing consortium from four candidate communities, each enriched for distinct SCFA profiles. This task required the integration of mechanistically grounded evidence retrieved through tool-mediated literature and database queries.

We next evaluated comparative therapeutic selection between two predefined synthetic consortia for DSS-induced colitis: an acetate-producing consortium (AceCom; *Dorea formicigenerans, Ruminococcus gnavus*, and *Blautia obeum*) and a lactate-producing consortium (LacCom; *Lactobacillus ruminis, Lactobacillus plantarum*, and *Streptococcus thermophilus*). The workflow comprised two phases:

- Phase I (Selection): Structured comparative evaluation integrating functional annotations, disease associations, and literature-derived mechanistic evidence to generate a ranked recommendation with supporting rationale.
- Phase II (Execution): Generation of an executable *in vivo* DSS colitis protocol specifying induction parameters, experimental groups, dosing schedules, and longitudinal inflammatory readouts.

For *de novo* consortium generation, Phase I was modified to constrain **Eubiota** to a laboratory inventory of 22 available type strains (Supplementary Note 6). Under this constraint, the system constructed a novel consortium by reasoning across domain-specific databases and literature tools, producing a mechanistically justified design. The selected community was subsequently passed to Phase II for protocol generation and experimental validation.

#### Mouse DSS colitis model validation

Colitis was induced in 8-week-old C57BL/6 wild-type mice (The Jackson Laboratory) using 2.5% (w/v) DSS (36–50 kDa, MP Biomedicals) administered *ad libitum* in autoclaved drinking water for 7 days, followed by a 3-day recovery period on regular water. Mice were age- and sex-matched and cohoused. For the AceCom versus LacCom comparison, experimental groups included: (i) water control, (ii) DSS only, and (iii) DSS supplemented with either AceCom or LacCom. For the *de novo* SynCom validation, groups included: (i) water control, (ii) DSS only, and (iii) DSS supplemented with the **Eubiota**-designed consortium. Individual strains were cultivated anaerobically for 48 h, pooled at equal volumes, and administered by oral gavage. Mice received daily gavage for 2 consecutive days prior to DSS exposure, followed by alternate-day dosing throughout DSS treatment and recovery. Disease progression was monitored daily via body weight and Disease Activity Index (DAI), scored based on weight loss, stool consistency, and rectal bleeding (QuantiQuik). Fecal lipocalin-2 was quantified by ELISA (R&D Systems). Colon length was measured at the experimental endpoint and analyzed using ImageJ.

### Antibiotic drug design targeting pathogens and pathobionts

#### Drug design workflow

A three-phase agentic workflow was implemented to design pathogen-biased antibiotic cocktails:

- Phase I (Cocktail Selection): Candidate antibiotic combinations were proposed by integrating pathogen susceptibility profiles, known resistance mechanisms, and drug-target pathway annotations.
- Phase II (Concentration Optimization): Minimum inhibitory concentration (MIC) data were aggregated across target pathogens and representative commensals to define concentration ranges that maximized inhibition of pathogens while minimizing commensal suppression.
- Phase III (Protocol Generation): An executable experimental protocol specifying dosing concentrations, growth conditions, and evaluation metrics was synthesized.

To improve selectivity, we incorporated a human-in-the-loop refinement cycle. Quantitative assay outputs (e.g., colony-forming unit (CFU) measurements) and qualitative observations were structured as feedback inputs, enabling **Eubiota** to iteratively adjust concentration ranges and recalibrate selective efficacy.

#### Antibiotic resistance experimental validation

To evaluate **Eubiota**-prioritized antibiotic cocktails, we selected two formulations corresponding to the upper and lower bounds of the predicted selective concentration window. Selective efficacy was assessed using high-density outgrowth assays against pathobionts (*P. aeruginosa* PA14, *C. portucalensis* HM-299, and *K. pneumoniae* NCTC 9633) and representative commensals (*E. coli* K12, *B. thetaiotaomicron* VPI-5482, and *B. breve* DSM 20213). All strains were grown to stationary phase in Brain Heart Infusion (BHI; BD Difco) medium. Cultures were diluted 1:2 into fresh BHI pre-supplemented with antibiotic cocktails and incubated statically at 37 °C for 24 h. Cocktail 1 contained amikacin (50 *µ*M), piperacillin (50 *µ*M), and tazobactam (13 *µ*M); Cocktail 2 contained amikacin (30 *µ*M), piperacillin (35 *µ*M), and tazobactam (13 *µ*M). Vehicle controls received sterile water.

Cell viability was quantified by CFU enumeration. Cultures were serially diluted in PBS, and 3 *µ*L aliquots were spotted onto BHI agar plates (ambient or pre-reduced, as appropriate), followed by aerobic or anaerobic incubation at 37 °C until countable colonies formed. The limit of detection was 333 CFU/mL. For experiments in which antibiotics were added at culture initiation, 48-h starter cultures were diluted 1:100 into fresh BHI containing the antibiotic cocktail and incubated for 48 h. Viability was then determined by CFU enumeration as described above.

### Novel anti-inflammatory molecule discovery from TWIN study

#### Anti-inflammatory molecule discovery workflow

We deployed **Eubiota** to prioritize candidate anti-inflammatory metabolites from a library of 397 circulating metabolites enriched under a vegan diet in the Twins Nutrition Study metabolomics dataset [131]. Metabolites were defined as vegan-enriched if they exhibited increased relative abundance compared with the omnivorous diet (log_2_ fold change of LC–MS peak intensity *>* 0). Detailed metabolomic acquisition and preprocessing procedures are described in the original study [131].

To navigate this high-dimensional search space, **Eubiota** integrated mechanistic relevance to inflammatory signaling with bibliometric novelty. Literature scarcity was quantified through iterative, tool-mediated retrieval, using the number and specificity of relevant hits as a proxy for prior characterization. Retrieved evidence was aggregated into a composite novelty score, with higher values reflecting limited representation in the biomedical literature.

To mitigate the stochasticity inherent in LLMs, each metabolite was evaluated across five independent workflow executions. In each run, metabolites were assigned a quantitative score (0–10) grounded in retrieved evidence, alongside categorical assessments of novelty (“highly novel” to “not novel”) and likelihood of anti-inflammatory activity (“highly likely” to “highly unlikely”). Final rankings were determined using the median rank across runs to enhance robustness against run-specific variance. In a subsequent phase, a Python-based module parsed and sorted the ranked outputs to identify the top 20 candidates. These high-priority metabolites were then passed to a final phase in which **Eubiota** synthesized detailed *in vitro* NF-*κ*B reporter assay validation protocols.

#### Tool usage and literature analysis

To characterize tool utilization, all computational and retrieval modules were enabled during execution. Comprehensive logs were recorded for each independent run, capturing tool invocations, input parameters, and returned outputs. Logs were parsed *post hoc* to extract and categorize tool-call events, enabling quantification of tool usage frequency and transition patterns across the workflow (Extended Data Fig. 4**c**). To assess evidence grounding, unique PubMed identifiers (PMIDs) referenced during execution were extracted from the system’s reasoning traces and enumerated. This analysis provided a quantitative measure of literature engagement and external evidence integration (Extended Data Fig. 4**d**).

#### Anti-inflammatory property validation via NF-*κ*B stimulation

Candidate compounds were dissolved in compatible solvents and diluted to working concentrations. A GFP-based NF-*κ*B reporter THP-1 cell line was used as previously described [132]. Cells were cultured in RPMI 1640 supplemented with 10% low-endotoxin fetal bovine serum (Thermo Fisher Scientific) and 1× penicillin-streptomycin at 37 °C in 5% CO_2_. Cells were harvested by centrifugation (500×*g*, 5 min), counted using trypan blue exclusion and a hemocytometer, and seeded into 96-well plates at 100,000 cells per well. After 24 h, cells were pre-treated with candidate compounds for 2 h prior to stimulation with either LPS (Sigma; 100 ng/mL) or Pam3CSK4 (InvivoGen; 100 ng/mL) for 18 h. Vehicle-only controls were included in all experiments.

NF-*κ*B activation was quantified by flow cytometry based on GFP fluorescence. Assay performance was confirmed by requiring that the mean fluorescence intensity (MFI) of stimulated positive controls exceeded unstimulated controls by at least 10-fold. Live cells were gated using forward scatter (FSC-A) and side scatter (SSC-A), followed by singlet discrimination using FSC-A/FSC-H (FlowJo). Experiments were repeated using independent clonal reporter lines to confirm reproducibility of NF-*κ*B activation responses.

## Author contributions

P.L. and Y.G. contributed equally to this work. P.L., Y.G., J.L.S., and J.Z. conceptualized the research. P.L. and Y.G. initiated and led the overall project. P.L., K.Z., H.Z., Q.X., B.L., W.G.P., and Y.G. developed the codebase and system. P.L., H.Z., and Q.X. contributed to data curation, system optimization, and benchmark evaluation. P.L., K.Z., H.Z., Q.X., and Y.G. developed the online interactive platform. Y.G., P.L., and W.G.P. developed scientific discovery tasks. Y.G., E.K.R., M.K., and A.L.S. conducted validation experiments. Y.G., P.L., W.G.P., E.K.R., and H.G.Z. analyzed validation results. Y.G. and P.L. led the human study. Y.C. and K.C.H. advised the project. J.L.S. and J.Z. supervised the project. All authors contributed to the preparation of the paper and approved the manuscript.

## Acknowledgements

We thank Bowen Chen, Sheng Liu, Fan Nie, Seungju Han, Fang Wu, Sherly Wu, Andrew Shen, Nitya Thakkar, Rubin Zou, Owen Queen, Zijian (Carl) Ma, Jake Silberg, Peter Eckmann, Alan Mao, Rahul Thapa, Sam Alber, and members of the Zou, Sonnenburg, Huang, and Choi groups for helpful discussions and feedback on this work. We also thank Kyle Swanson, Wanjia Zhao, Mert Yuksekgonul, Haotian Ye, Aneesh Pappu, Elana Simon, Batu El, Mirac Suzgun, Joseph Boen, Arushi Gupta, Siyu He, and Jiacheng Miao for their support. We are grateful to Tony Xia for his assistance with system prototype development and acknowledge Weston Whitaker, Achuthan Ambat, and Sean Spencer for early project discussions. We appreciate the generous help of Mikhail Iakiviak from Microbiome Therapies Initiative (MITI) for providing bacterial type strains. We also sincerely thank Matthew Carter and the Christopher Gardner group at Stanford School of Medicine for sharing data from the TWIN study and for foundational work establishing the cohorts. We acknowledge the help from Steven Higginbottom for the mouse work. We thank our human study volunteers, including Erica Sonnenburg, Anita Reddy, Tadashi Takeuchi, Ian Miller, Sriteja Kataru, Samantha Goldman-Hernandez, Sailendharan Sudakaran, Maggie Madrigal-Moeller, and anonymous participants for their time and thoughtful feedback. This work was supported by funding from Stanford HAI and the AI for Math Fund by Renaissance Philanthropy, and the NIH (R01DK085025 to J.L.S.). J.Z. and J.L.S. are Chan Zuckerberg Biohub Investigators.

## Data availability

All data used to train and evaluate **Eubiota** are available in the project repository at https://github.com/lupantech/Eubiota. This includes the processed training and benchmark datasets, derived from publicly available and expert-curated sources, along with documentation and instructions for reproduction.

## Code availability

Source code for **Eubiota** is available at https://github.com/lupantech/Eubiota. To facilitate access and exploration, a live interactive platform is available at: https://app.eubiota.ai, and an online research workflow interface is provided at https://app.eubiota.ai/workflow.

## Extended Data

**Extended Data Figure 1:**
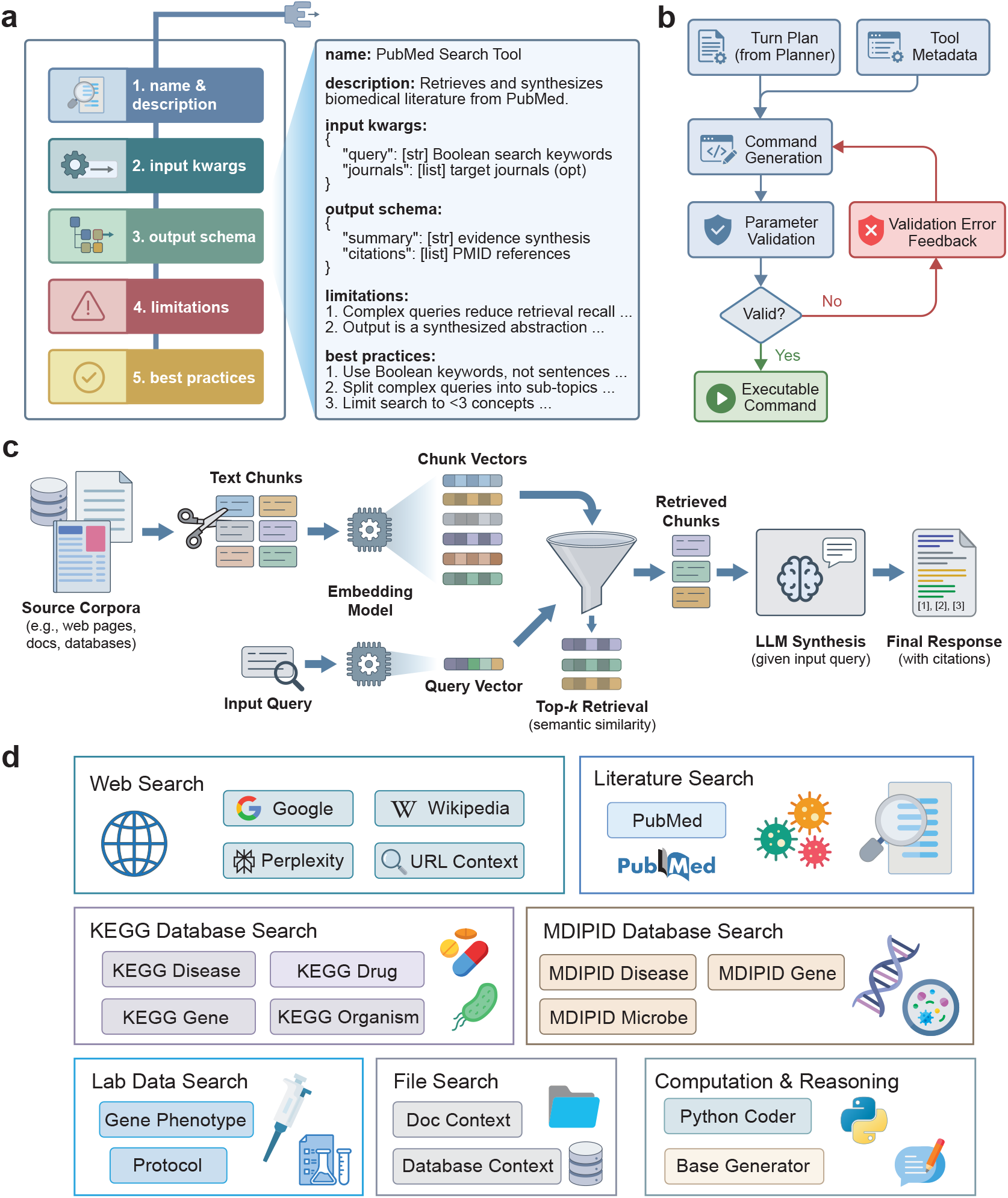
Unified tool interface and domain-specific ecosystem. **a**, Unified tool interface specification enabling rapid definition and seamless routing of custom tools. **b**, Command generation process with self-refinement to ensure parameter validity. **c**, Retrieval-augmented generation (RAG) mechanism for knowledge tools. Source documents are chunked, embedded, and retrieved based on vector similarity to ground LLM generation in factual evidence. **d**, Domain-specific toolset comprising modules for laboratory document retrieval, web and literature search, and structured biological databases (KEGG, MDIPID).

**Extended Data Figure 2:**
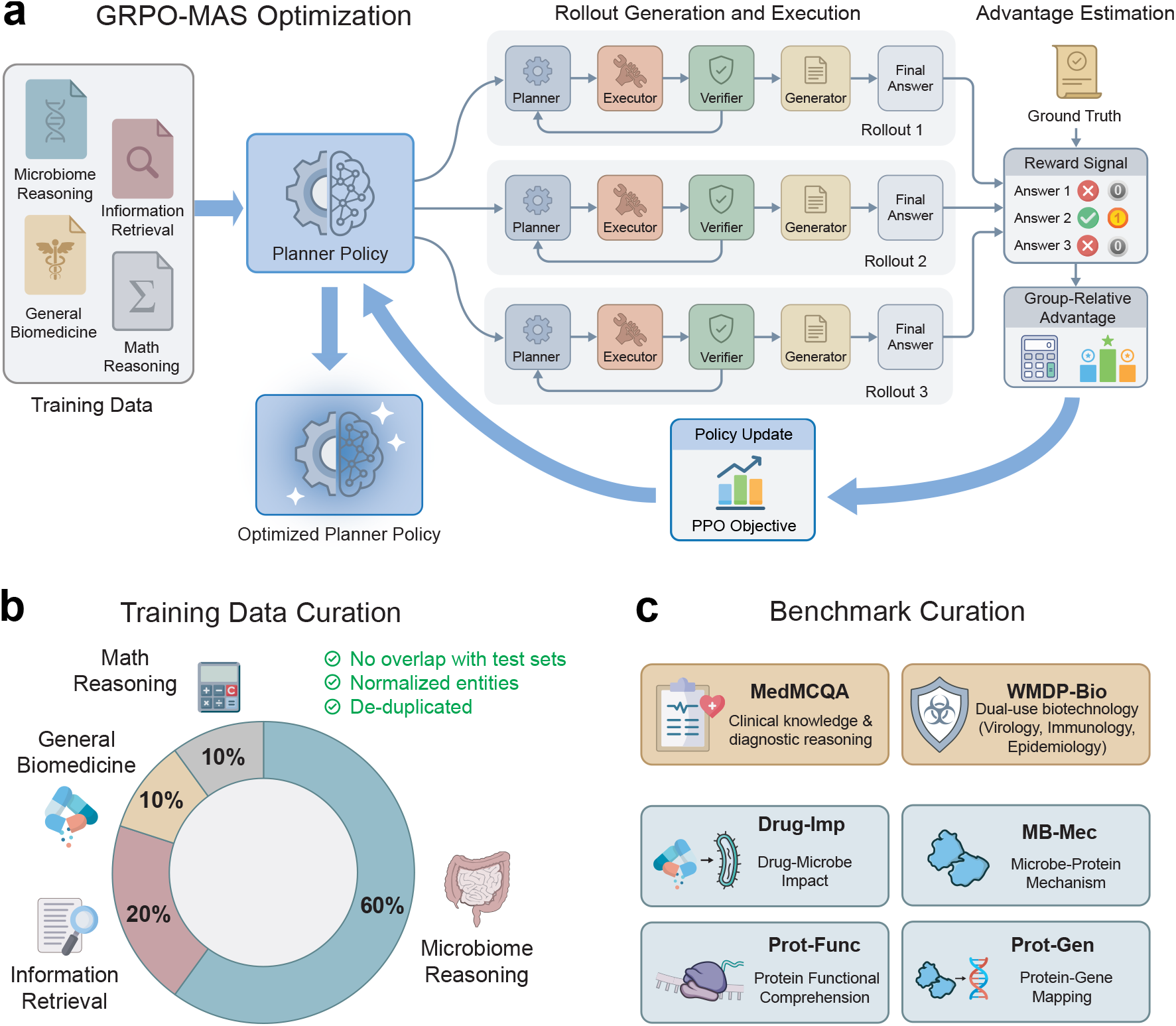
System optimization and evaluation. **a**, Schematic of the GRPO-MAS optimization strategy used to train the Planner policy prior to deployment. During training, the system generates a batch of independent multi-turn rollouts via stochastic action sampling (temperature = 0.7), enabling the exploration of diverse reasoning trajectories for the same input query. Outcomes are evaluated against ground truth to compute group-relative advantages, which drive the policy update to reinforce successful reasoning traces. Outcomes are evaluated against ground truth to update the policy; the resulting Optimized Planner Policy is then frozen and integrated into the core inference framework (Fig. 1**a**). **b**, Composition of the training dataset (*N* = 2,000) stratified by domain: microbiome reasoning (60%), information retrieval (20%), general biomedicine (10%), and math reasoning (10%). Data were curated via deduplication, entity normalization, and strict removal of overlap with evaluation benchmarks. **c**, Benchmark evaluation suite comprising general biomedical assessments (MedMCQA Medicine subset and WMDP-Bio) and four domain-specific mechanistic reasoning tasks derived from the MDIPID database: drug–microbe impact (Drug-Imp), microbe–protein mechanism (MB-Mec), protein functional comprehension (Prot-Func), and protein–gene mapping (Prot-Gen).

**Extended Data Figure 3:**
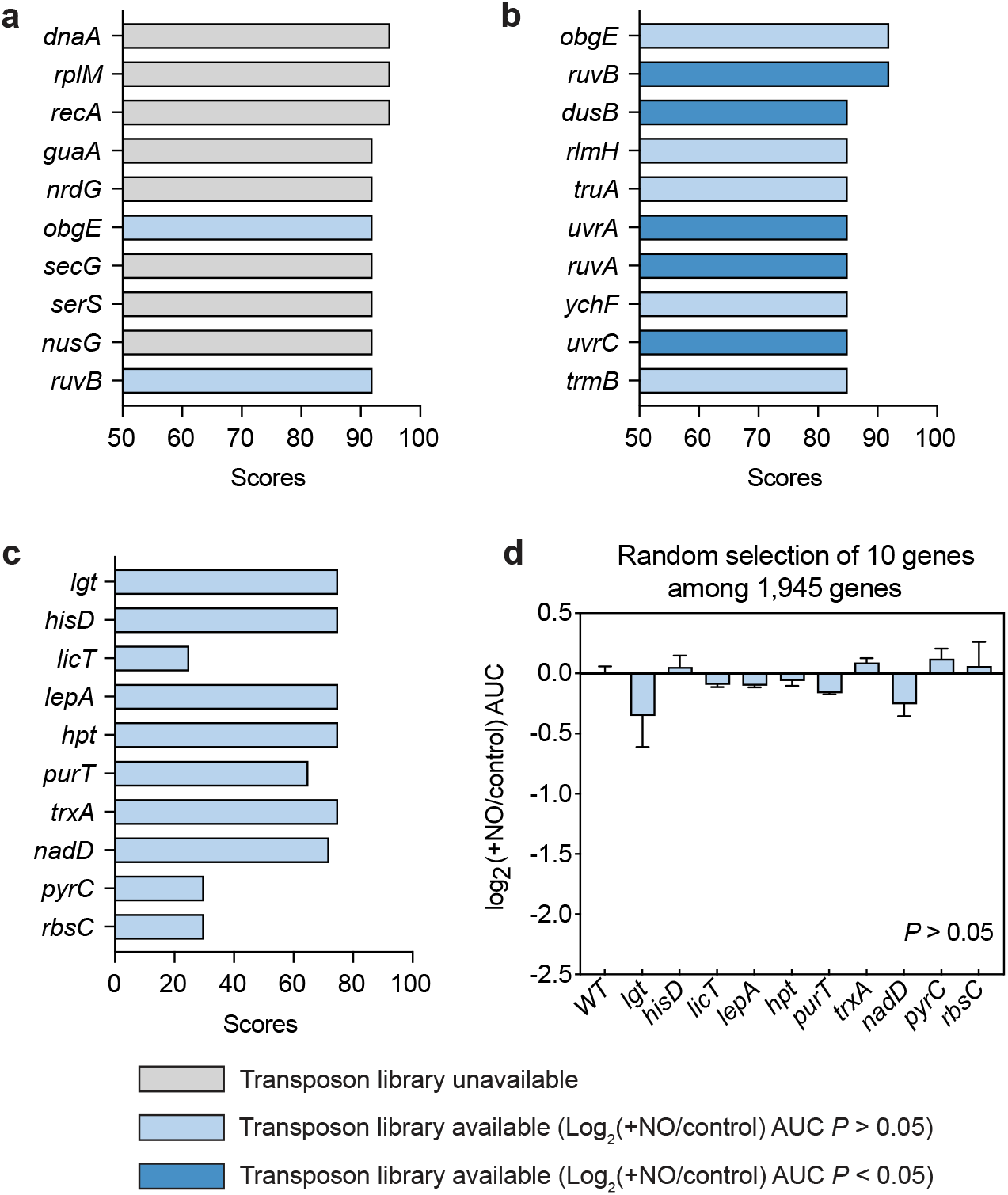
Experimental validation of prioritized genes. **a**, The top 10 genes prioritized by **Eubiota** in the full list of 1,945 genes. **b**, The top 10 genes with transposon mutant available prioritized by **Eubiota** in the full list of 1,945 genes. **c**, Ten genes randomly selected from the full list of 1,945 genes. **d**, The growth of randomly selected 10 genes from the 1,945 global list. Plotted is the area under the curve (AUC) change log_2_(AUC_NO_/AUC_control_) (*n* = 3). No significant growth inhibition was observed. Values are represented as mean ± s.e.m., and *P* values were calculated using two-tailed Student’s *t*-test on raw AUC. Grey color represents transposon mutants that are not available. Light blue represents transposon mutants available for experimental validation and the AUC change is not significant (*P >* 0.05). Dark blue represents transposon mutants available for experimental validation and the AUC change is significant (*P* < 0.05)

**Extended Data Figure 4:**
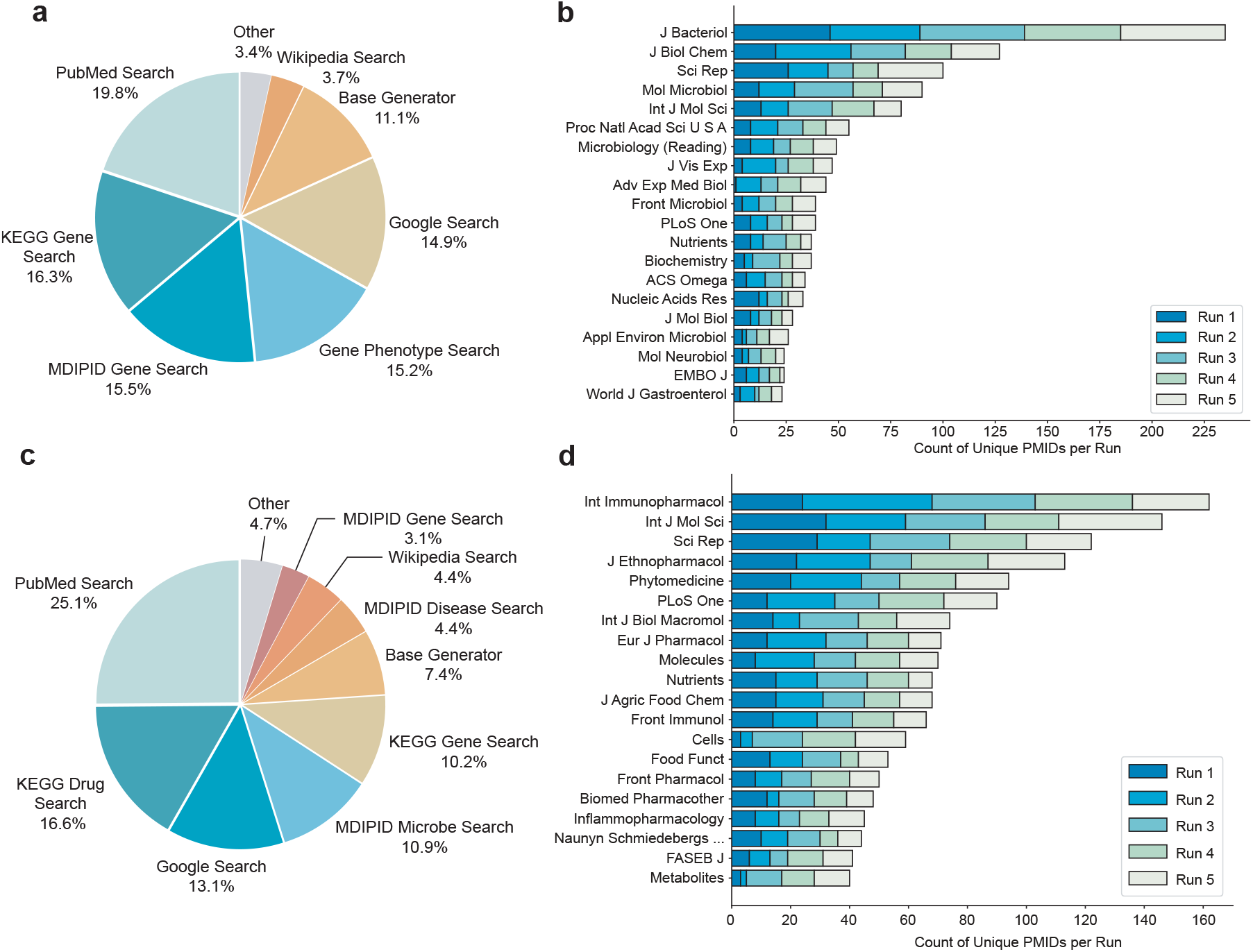
Literature and tool use metrics across genetic discovery and metabolite discovery tasks. **a**, Tool usage distribution pie chart in genetic discovery. Across five runs, PubMed Search, KEGG Gene Search, MDIPID Gene Search, Gene Phenotype Search, and Google Search were the most-used tools. Tool calls were concentrated in task-relevant modules across runs. **b**, Analyses of journals cited across runs in genetic discovery. Top-cited journals are relevant to bacteriology, microbiology, biochemistry, and nutrition, aligning with the genetic discovery task. **c**, Tool usage distribution pie chart in molecular discovery. Across five runs, PubMed Search, KEGG Drug Search, Google Search, MDIPID Microbe Search, and KEGG Gene Search were the most-used tools. Calls to irrelevant tools were significantly decreased. **d**, Analyses of journals cited across runs in molecular discovery. Top-cited journals are relevant to immunology, pharmacology, nutrition, and metabolites, aligning with the objective of NF-*κ*B inhibition.

**Extended Data Figure 5:**
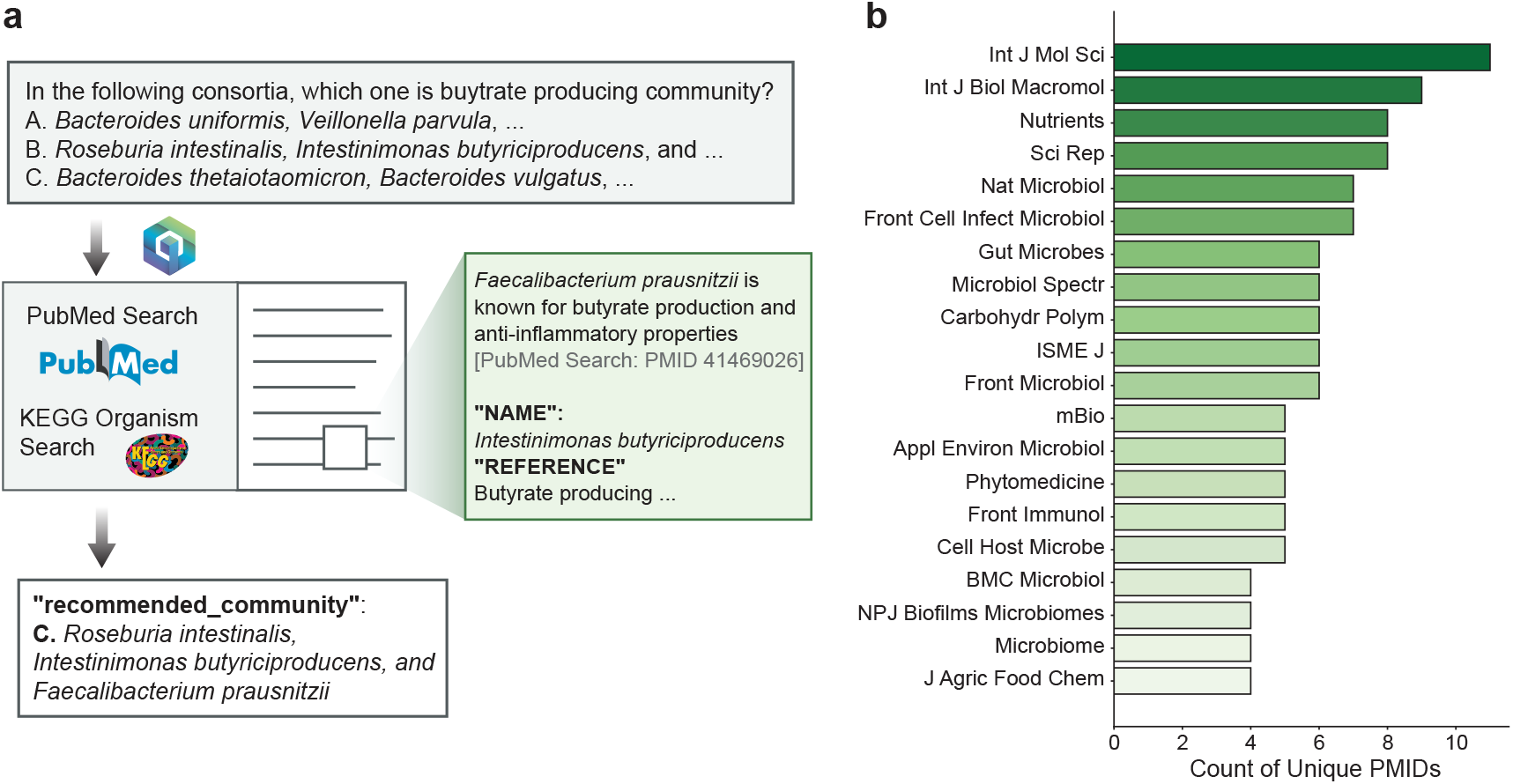
Synthetic microbial community for gut diseases task. **a**, A query was given to **Eubiota** to pick a known butyrate-producing consortium from three options. The test was run three times, with the correct consortium shuffled among choices A, B, and C. The system correctly identified the community each time, citing specific literature-based evidence for its selection. **b**, Top 20 journals by citation frequency.

**Extended Data Figure 6:**
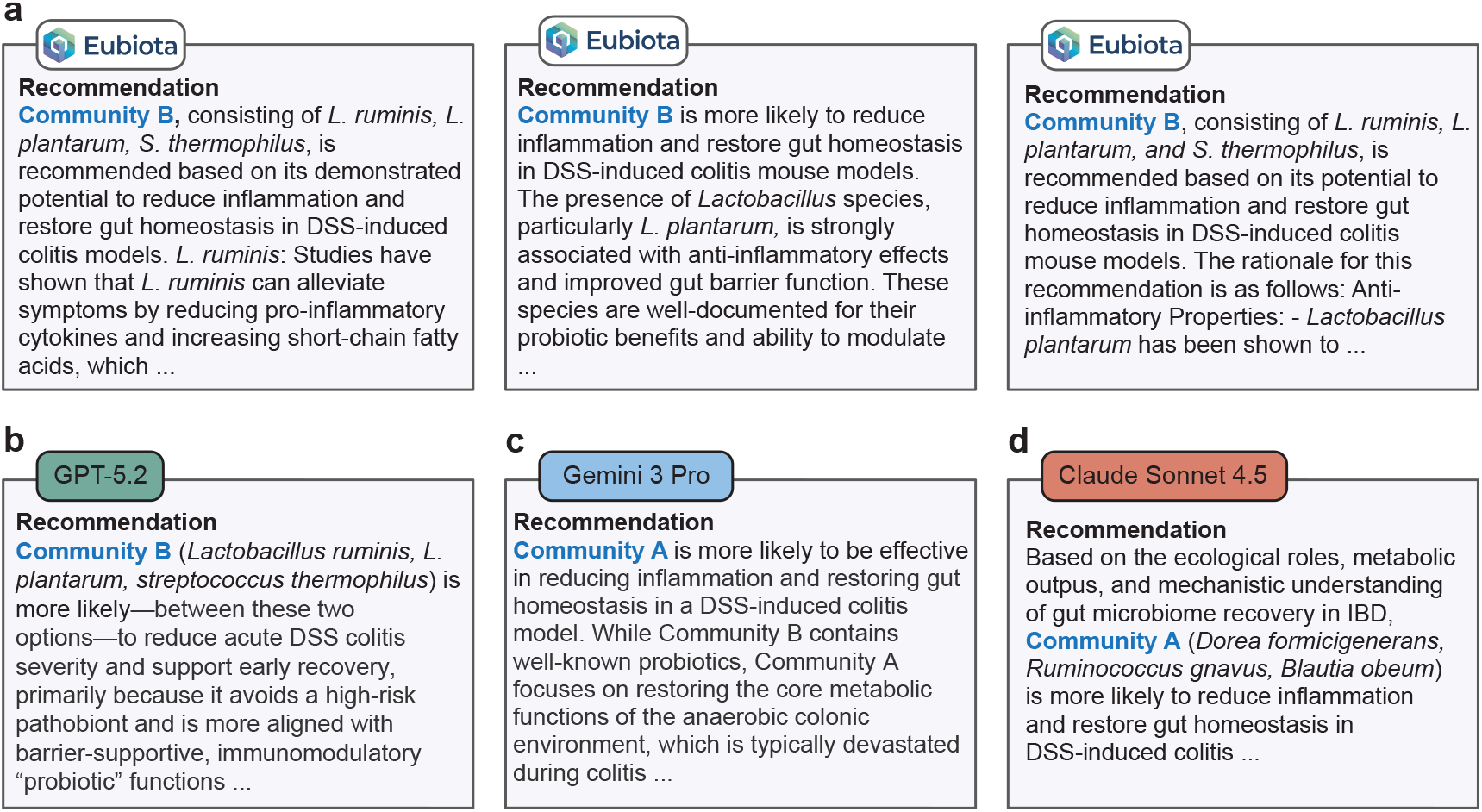
Comparative reasoning against frontier models. **a**, Three runs of **Eubiota**, showing consistent selection. **b-c**, Three frontier models (GPT-5.2 [106], Gemini 3 Pro [107], and Claude Sonnet 4.5 [108]) were tested with the same query, yielding differing recommendations and rationales.

**Extended Data Figure 7:**
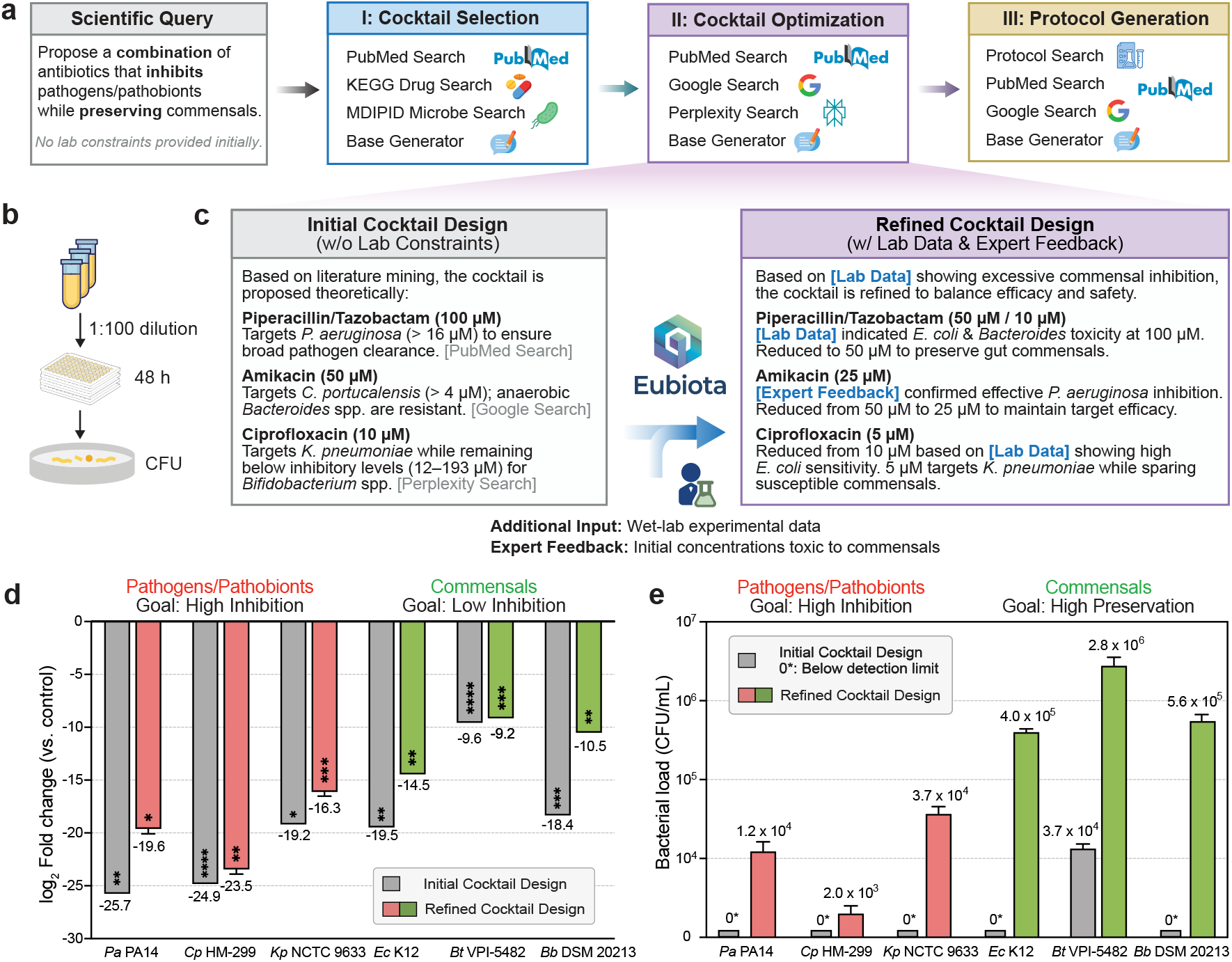
Interactive feedback configuration of Eubiota guided antibiotic cocktail design. **a, Eubiota** pipeline consists of three phases: Phase I (Cocktail Selection) identifies candidate antibiotics via PubMed, KEGG Drug, and MDIPID Microbe Search; Phase II (Cocktail Optimization) refines these selections using broad-scale search engines and generative models; and Phase III (Protocol Generation) produces standardized laboratory instructions. **b**, Experimental validation workflow. Bacterial cultures are diluted 1:100 and incubated anaerobically for 48 hours. Viability is subsequently assessed via colony-forming unit (CFU) quantification. **c**, Iterative refinement of initial antibiotic cocktail design. By integrating expert feedback, the concentrations were significantly reduced to achieve a more precise formulation. **d**, The differential bacterial suppression of the initial (grey) and refined (colored) antibiotic cocktail designs were compared across a panel of pathogens/pathobionts (*P. aeruginosa* PA14, *K. pneumoniae* NCTC 9633, *C. portucalensis* HM-299) and commensals (green: *E. coli* K12, *B. thetaiotaomicron* VPI-5482, and *B. breve* DSM 20213), shown as the log_2_ change in CFU/mL. While both designs exhibit inhibition of pathobionts (red), the refined cocktail reduces off-target toxicity toward beneficial commensal species (green), such as *Bt* VPI-5482 and *Bb* DSM 20213. **e**, Absolute quantification of bacterial load (CFU/mL) following treatment, demonstrating that the refined design maintains high inhibition against pathobionts while preserving commensals relatively. 0* denotes below limit of detection (LOD = 333 CFU/mL). Data are presented as mean ± s.e.m. *P* values were determined by twotailed Student’s *t*-test on raw CFU values to compare mean differences between antibiotic-treated and untreated conditions. ^∗^*P* < 0.05, ^∗∗^*P* < 0.01, ^∗∗∗^*P* < 0.001, ^∗∗∗∗^*P* < 0.0001.

**Extended Data Table 1:** Efficiency comparison between expert curation and Eubiota. Each run reports the number of unique PubMed articles processed, estimated human expert time, actual **Eubiota** runtime, and resulting fold efficiency gain. To estimate expert time consumption, we assume an expert takes 3 minutes to review one paper. **Eubiota** experiments were conducted on two NVIDIA A100 GPUs; efficiency can scale with additional compute resources.

| Run | Unique PMIDs | Est. expert time (h) | Eubiota run time (h) | Fold efficiency |
| --- | --- | --- | --- | --- |
| 1 | 804 | 40.20 | 1.24 | ×32.4 |
| 2 | 969 | 48.45 | 1.13 | ×42.9 |
| 3 | 945 | 47.25 | 1.14 | ×41.5 |
| 4 | 850 | 42.50 | 1.35 | ×31.5 |
| 5 | 1,033 | 51.65 | 1.14 | ×45.3 |
| **Mean** | **920.2** | **46.01** | **1.20** | **×38.7** |

## Supplementary Information

### Supplementary Note 1: Eubiota usage modes and user interface

To bridge the gap between advanced computational reasoning and routine experimental practice, **Eubiota** is offered through flexible operating modes that support inquiry at multiple scales, ranging from single-query exploration to configuration-driven, high-throughput pipeline execution (Supplementary Fig. 1).

**Supplementary Figure 1:**
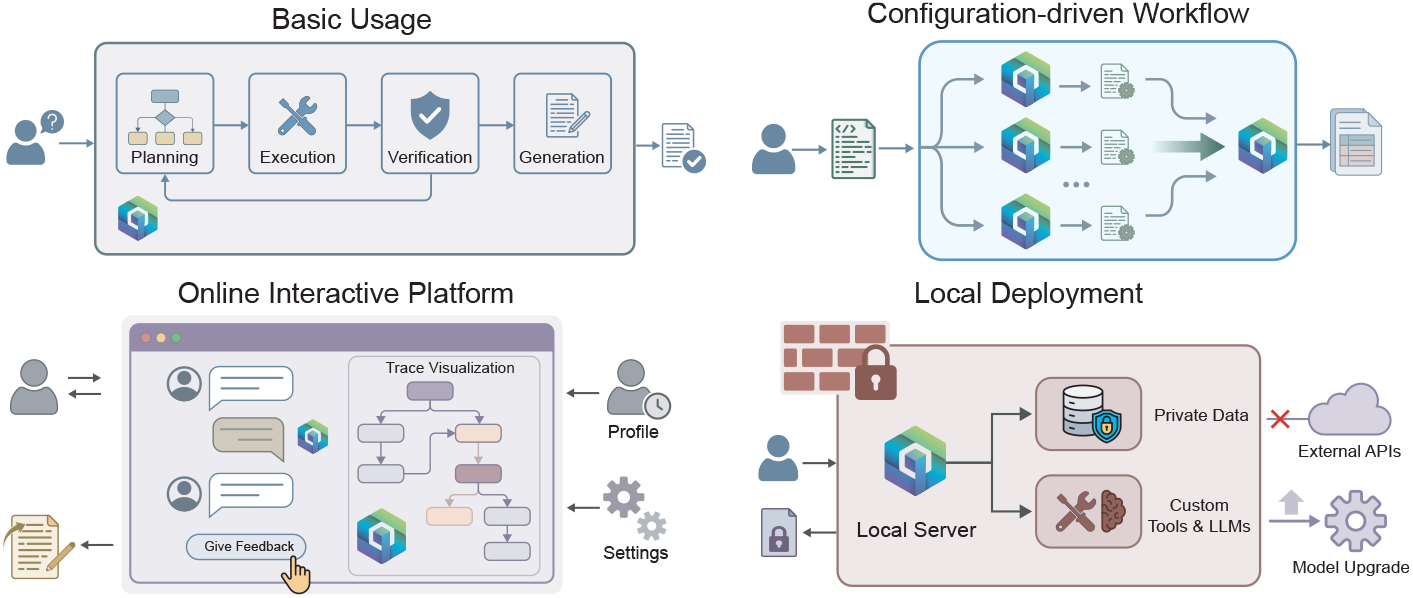
Overview of Eubiota usage modes. The framework supports a spectrum of needs from basic interactive queries to large-scale, secure discovery workflows.

#### Basic usage mode

In its basic mode, **Eubiota** processes individual scientific queries through an iterative cycle of planning, execution, verification, and grounded generation (Main Text 2.3). Intermediate results, reasoning traces, and retrieved evidence are recorded in shared memory, ensuring that each response is grounded in the execution history.

#### Configuration-based discovery workflows

For large-scale discovery tasks, **Eubiota** supports a configuration-driven mode in which researchers specify workflow constraints across phases using structured configuration files. Each phase defines subgoals, batch inputs, and formatted outputs. For example, in the gene prioritization task (Main Text 3.2), an initial phase screens genes in parallel and passes ranked candidates to downstream phases for experimental protocol generation. This design fixes workflow specifications to enable reproducible execution of parallelized reasoning trajectories over thousands of candidates without manual intervention during runtime.

#### Online interactive platform

**Eubiota** is accessible through a web interface that supports multiturn natural language interactions and visual monitoring of structured workflows (Supplementary Figs. 2 and 3). The interface enables users to inspect reasoning traces, review intermediate tool outputs, and provide corrective feedback in real time, thereby maintaining human oversight throughout the decision process. The platform also maintains chat history and user-specific settings to support contextual continuity across sessions.

#### Local deployment and extensibility

To address the data privacy requirements of clinical research, we provide an open-source implementation that supports local deployment behind institutional firewalls, with external API calls disabled when required by policy. The modular design further allows upgrading the LLM backbone or substituting open-weights models fine-tuned on institution-specific data, enabling adaptation to new tool ecosystems and biological domains.

**Supplementary Figure 2:**
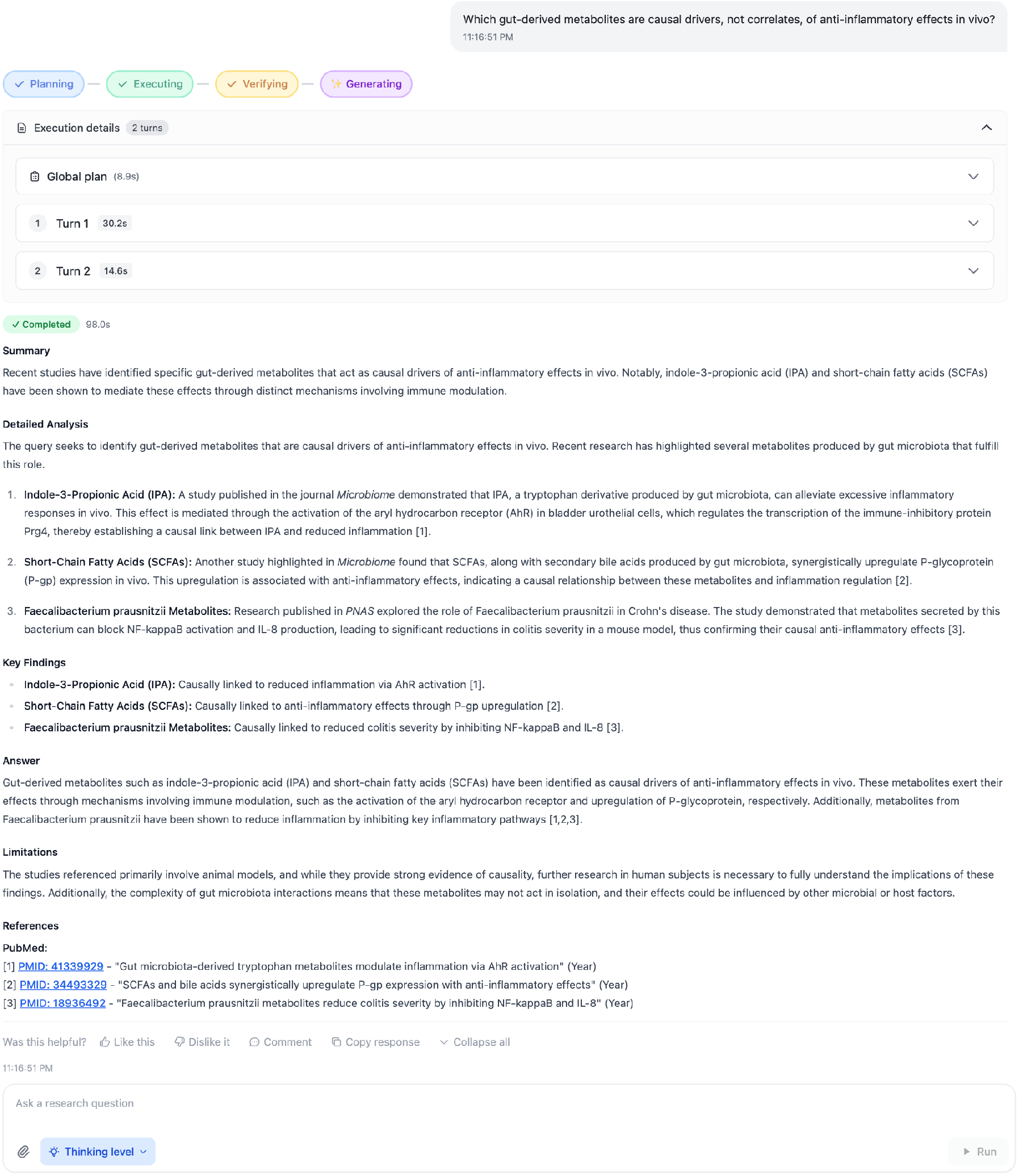
The Eubiota interactive chat interface. Researchers input research questions and configure inference parameters, such as reasoning depth, prior to execution. The system visualizes the Planning, Executing, Verifying, and Generating steps in real time. Through multi-turn iterations, **Eubiota** synthesizes comprehensive answers supported by valid citations.

**Supplementary Figure 3:**
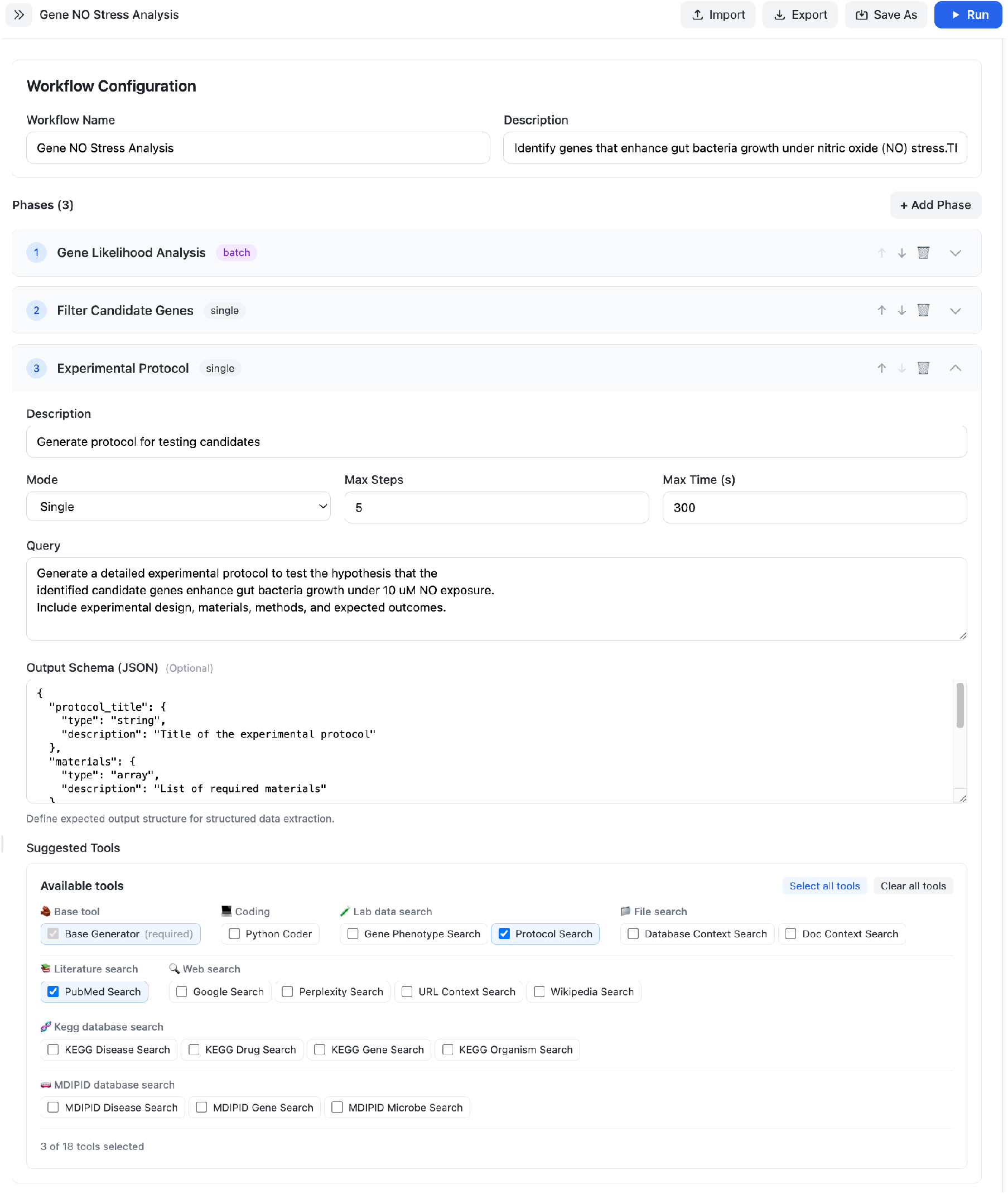
Configuration-based discovery workflows in Eubiota. Researchers design structured experimental pipelines to drive real-world discovery. Users define distinct phases within the experiment and customize tool selection for each stage. Workflows can be imported, exported, and saved to the library to ensure reproducibility.

### Supplementary Note 2: System design and inference logic

Extended Data Fig. 2 illustrates the high-level architecture of the **Eubiota** framework. Here, we present the formal inference procedure (Algorithm 1) that specifies how the Planner, Executor, Verifier, and Generator coordinate over multiple turns through a shared, evolving memory state ℳ_*t*_. At each turn *t*, the Planner proposes a subgoal and selects candidate tools based on ℳ_*t*_; the Executor issues validated tool calls and records observations; and the Verifier evaluates whether the accumulated evidence satisfies the stopping criteria, updating ℳ_*t*+1_. Once termination is triggered, the Generator synthesizes the final evidence-grounded response from the full memory trace.

#### Algorithm 1

**Eubiota** modular agentic inference process

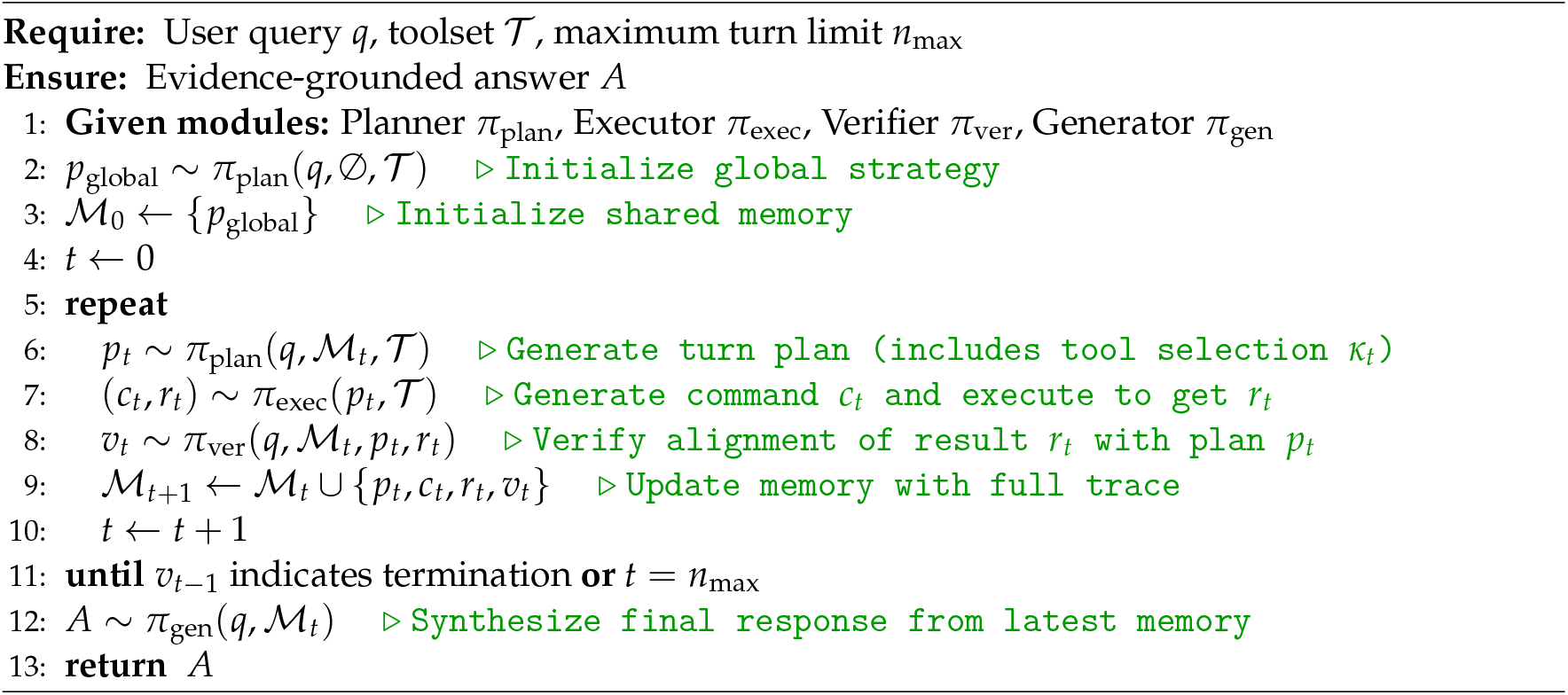

### Supplementary Note 3: Tool ecosystem and interface specifications

To enable reliable and extensible tool use, **Eubiota** implements a standardized tool interface that treats each external capability as a typed, auditable module. Each tool is defined by (i) a unique name and description, (ii) typed input arguments with validation rules, (iii) a structured output schema, and (iv) documented limitations and best practices that constrain the Planner to produce executable calls. This design decouples agent reasoning from tool implementations, allowing new tools to be added or swapped without modifying the core multi-agent logic, while ensuring that intermediate evidence (queries, retrieved records, and tool outputs) is consistently logged into shared memory for downstream verification and traceability. Extended Data Fig. 1**a** illustrates the unified tool specification and routing mechanism, and Extended Data Figs. 1**c** and 1**d** summarize the retrieval-backed tools and the domain-specific tool ecosystem used in this study.

To illustrate the standardized interaction between the Executor and the domain-specific toolset, we provide detailed specifications and interaction examples for four representative tools. The tool definitions presented here have been slightly simplified and reformatted for clarity while maintaining the authentic logic of the codebase. Each tool defines specific limitations and best practices to guide the Planner in generating valid and efficient execution strategies.

#### PubMed Search

This tool serves as the primary gateway for retrieving biomedical and life sciences literature from the PubMed database. It is designed to synthesize abstracts into evidence-grounded summaries with PMID-traceable citations, leveraging Boolean logic (using AND to combine keywords) to maximize retrieval precision and minimize extraneous noise.

##### PubMed Search Tool: Metadata

**Description**: A comprehensive tool for searching the PubMed database, the primary database for biomedical and life sciences literature.

**Limitations**:

1. Limited to the PubMed database.
2. Extremely specific combinations (e.g., long lists of species/strains/genes plus model plus treatment) may yield no results. Split these into individual queries.
3. Results may include non-peer-reviewed items; always verify relevance and quality.

**Best practices**:

1. Suited for searching general information on biomedical topics.
2. Do not enumerate many species or genes. Keep queries under 3 topics.
3. Use bag-of-words rather than sentences. Combine keywords with “AND”.
4. For complex questions (e.g., condition + model + intervention), search each axis separately and combine downstream.
5. Scan abstracts post-retrieval to confirm biological context (organism, tissue, assay) and relevance.

##### PubMed Search Tool: Usage example

~~~
**Tool calling arguments:**
{
 “query”: “CRISPR gene editing applications”,
 “journals”: [“Nature”, “Science”],
 “max_results”: 5
}
**Final output from the Executor:**
{
 “Nature”: {
  “term_used”: “CRISPR gene editing applications AND Nature[Journal]”,
  “summary”: “Recent publications in Nature discuss the therapeutic…”,
  “citations”: [
   “[1] PMID: 38123456”,
   “[2] PMID: 39123457”
  ]
 },
 “Science”: {
  “term_used”: “CRISPR gene editing applications AND Science[Journal]”,
  “summary”: “Science reports focus on off-target effects…”,
  “citations”: [“…”]
 }
}
~~~

#### KEGG Drug Search

This tool provides access to structured pharmacological knowledge within the local KEGG database. Unlike general search tools, it returns raw JSON data fields (e.g., efficacy, formula), making it ideal for precise fact-checking but requiring exact or high-similarity drug name matches.

##### KEGG Drug Search Tool: Metadata

**Description**: A specialized tool for searching the KEGG database to find information about a drug (e.g., category, efficacy, etc.) given a drug name.

**Limitations**:

1. Can only return biomedical domain knowledge given a drug name or keyword.
2. Returns raw JSON data rather than a summarized narrative answer.

**Best practices**:

1. Use for structured drug facts (e.g., entry ID, formula, efficacy) from the local cache.
2. Query with specific identifiers: exact generic name, brand name, or known synonym.
3. Keep queries short and name-like. Semantic queries are allowed but result in approximate matches.
4. Results are ranked by embedding similarity. Always confirm the returned “NAME” matches the intended drug.
5. Run one query per drug; do not combine multiple drugs into a single query.

##### KEGG Drug Search Tool: Usage example

~~~
**Tool calling arguments:**
{
 “query”: “Aspirin”,
 “keys_to_output”: [“ENTRY”, “NAME”, “FORMULA”, “EFFICACY”],
 “max_results”: 1
}
**Final output from the Executor:**
{
 “status”: “Success”,
 “results”: [
  {
    “data”: {
      “ENTRY”: “D00109”,
      “NAME”: “Aspirin (USP); Acetylsalicylic acid (JP17)”,
      “FORMULA”: “C9H8O4”,
      “EFFICACY”: “Analgesic, Anti-inflammatory, Antipyretic”
    },
    “similarity”: 1.0
  }
 ]
}
~~~

#### MDIPID Microbe Search

Designed for complex microbiome inquiries, this tool combines database retrieval with LLM synthesis. It addresses the limitation of pure retrieval by generating a natural language answer to a specific question while providing truncated raw data as verifiable evidence.

##### MDIPID Microbe Search Tool: Metadata

**Description**: A specialized tool for searching the MDIPID database. It returns a concise answer and truncated structured data as evidence. Useful for taxonomy, drug interactions, and disease associations.

**Limitations**:

1. Can only search by microbe name.
2. Returns distinct `llm_answer` and truncated `evidence_data`.
3. `evidence_data` is strictly truncated; do not assume it is the complete list.
4. Requires specific `info_to_search` question and `query` (taxon name).

**Best practices**:

1. `query` must be one specific microbe name (e.g., “Phocaeicola vulgatus”).
2. Always select specific `keys_to_output` (e.g., “microbiota-disease associations”) to prevent context overflow.
3. Phrase `info_to_search` as a clear, standalone question.
4. Prioritize information in `llm_answer` as it saw the untruncated data.

##### MDIPID Microbe Search Tool: Usage example

~~~
**Tool calling arguments:**
{
 “query”: “Phocaeicola vulgatus”,
 “info_to_search”: “What diseases is this microbe associated with?”,
 “keys_to_output”: [“microbiota-disease associations”]
}
**Final output from the Executor:**
{
 “status”: “Success”,
 “results”: [
   {
     “microbe_id”: “MIC00123”,
     “match_type”: “exact”,
     “llm_answer”: “Phocaeicola vulgatus is positively associated with…”,
     “evidence_data”: “{
     “microbiota-disease associations”: [
       {“disease”: “IBS”, “association”: “positive”}
       ]
     }”
   }
 ]
}
~~~

#### Google Search

To bridge the gap between static databases and real-time knowledge, **Eubiota** integrates a Google Search tool. This module is optimized for general information retrieval and automatically enriches results with markdown-formatted citations to ensure traceability.

##### Google Search Tool: Metadata

**Description**: A web search tool powered by Google’s Gemini AI that provides real-time information from the internet with citation support.

**Limitations**:

1. Only suitable for general information search.
2. Contains less domain-specific information.
3. Not suitable for searching or analyzing videos.

**Best practices**:

1. Use for definitions, world knowledge, and general information.
2. Query must be specific, clear, and self-contained (avoid vague references like this “mentioned earlier”).
3. Explicitly include full names/identifiers; do not assume the tool has access to context from previous steps.
4. Returns summarized information with citations.

##### Google Search Tool: Usage example

~~~
**Tool calling arguments:**
{
   “query”: “What are the mechanism of action for Metformin?”,
   “add_citations”: true
}
**Final output from the Executor:**
{
   “response”:
      “Metformin, a frontline medication for type 2 diabetes, employs a complex and
      multifaceted mechanism of action to lower blood glucose levels.
      Its effects primarily target the liver and the gut, while also influencing
      systemic insulin sensitivity. Key mechanisms of action for Metformin include:
      **Inhibition of Hepatic Glucose Production:** Metformin’s primary action
      involves reducing the liver’s output of glucose. It achieves this by inhibiting
      Complex I of the mitochondrial respiratory chain in liver cells
      [1] (https://pmc.ncbi.nlm.nih.gov/articles/PMC5552828/),
      [2] (https://en.wikipedia.org/wiki/Metformin),
      [3] (https://diabetesjournals.org/care/article/39/2/187/37215/Mechanism-of-Metformin-A-Tale-of-Two-Sites).
      This action …”
}
~~~

### Supplementary Note 4: Agent system prompts and shared memory schema

The **Eubiota** framework uses structured, role-specific system prompts to coordinate four specialized agents: Planner, Executor, Verifier, and Generator. We present the core instructions from the system implementation and the structured Memory format used for inter-agent communication. Memory stores the global strategy and a turn-indexed log of execution steps. Each step records the states of the Planner, Executor, and Verifier; the Generator is invoked only after the Verifier terminates the loop to synthesize and record the final, evidence-grounded response.

#### Planner Agent: System prompt

▽ *Initial, global planning phase*

**Objective:** Analyze the query and create an execution plan.

**Input context:** Query, available tools, tool metadata, files.

**Required output:**

1. Decomposition of query objectives
2. Required skills
3. Relevant tools with justification
4. Additional considerations

▽ *Turn planning phase*

**Objective:** Determine the optimal next step to address the query using available tools and previous steps.

**Input context:** Query, global plan, tools, metadata, files, previous steps, turn budget.

**Operational instructions:**

1. Analyze the query, plan, and history to identify what remains.
2. Review tool metadata (limitations, best practices).
3. Select the optimal tool (single selection) to maximize progress.
4. Formulate a specific, achievable subgoal.
5. Specify all necessary context (variables, data flow).
6. Avoid redundancy by building on previous results.

**Required output schema:**

1. Step analysis: Justification for tool choice and approach.
2. Context: Complete file paths, variable names, and data values.
3. Subgoal: Specific objective for this turn.
4. Tool name: Exact registry match (no variations).
5. Specify all necessary context (variables, data flow).
6. Expected outcome: Predicted result of the turn.

#### Executor Agent: System prompt

**Objective:** Translate the planned action into precise, executable tool arguments.

**Input context:** Subgoal, context variables, selected tool definition, tool schema.

**Operational instructions:**

1. Review the tool’s `input_kwargs` schema for parameter requirements.
2. Generate JSON commands using exact, case-sensitive parameter names.
3. Enforce strict data typing (string, boolean, integer, array, object).
4. Include all required parameters; omit optional ones if unnecessary.
5. Format output as a single JSON object or a list of objects for batching.
6. Critical: Never use placeholder values.

**Required output schema:**

1. Analysis: Brief logic for parameter construction.
2. Generated arguments: Valid JSON block (e.g., `“param”:”value”`).

#### Verifier Agent: System prompt

**Objective:** Assess the sufficiency of accumulated evidence to resolve the query.

**Input context:** Query, global plan, tool definitions, files, memory state.

**Evaluation criteria:**

1. Completeness: Have all aspects of the global plan been addressed?
2. Tool utility: Can remaining tools provide critical missing evidence?
3. Consistency: Are there contradictions within the retrieved data?
4. Verification: Do specific claims require independent cross-checking?
5. Ambiguity: Are current results sufficiently precise?

**Required output schema:**

1. Analysis: Detailed evaluation addressing the criteria above.
2. Conclusion: Binary state (`“STOP”` or `“CONTINUE”`) to control loop termination.

#### Generator Agent: System prompt

**Objective:** Synthesize a grounded, comprehensive scientific response based on the execution trace.

**Input context:** Query, global plan, file context, full execution history.

**Required output structure:**

1. Executive summary: Brief overview of the research goal and outcome.
2. Methodological walkthrough: Detailed narrative of tool usage, key intermediate results, and evidence integration.
3. Key findings: Specific biological discoveries or data points extracted.
4. Direct response: Clear, explicit answer to the initial query.
5. Critical assessment: Identification of limitations, data gaps, or caveats.
6. Conclusion: Final synthesis and recommendations for next steps.

#### Shared memory schema (JSON structure)

~~~
{
  “global_plan”: {
    “strategy”: “1. Search literature for gene X… 2. Analyze phenotypes…”,
    “execution_time”: 12.5
    },
  “turn_history”: {
    “Turn 1”: {
      “Planner”: {
        “tool_to_use”: “PubMed_Search”,
        “turn_goal”: “Retrieve papers on recR and NO stress”,
        “turn_context”: “Query: recR AND nitric oxide…”,
        “execution_time”: 8.5
      },
      “Executor”: {
        “generation_analysis”: “Constructing query for PubMed…”,
        “generation_time”: 5.0,
        “execution_time”: 2.1,
        “execution_results”: [
          {
             “tool_name”: “PubMed_Search”,
             “command_arguments”: { “query”: “recR nitric oxide” },
             “summary”: “Found 5 relevant papers…”,
             “publications”: […]
          }
        ]
      },
      “Verifier”: {
        “analysis”: “Evidence found supports the hypothesis but lacks…”,
        “stop_signal”: false,
        “execution_time”: 6.0
      }
    },
    “Turn 2”: { … }
  }
}
~~~

### Supplementary Note 5: Training data curation and system evaluation

#### Training data curation

As summarized in the main text and Extended Data Fig. 2**b**, we curated a 2,000-instance training corpus with a fixed 6:2:1:1 mixture across microbiome reasoning, information retrieval, general biomedical knowledge, and mathematical reasoning. Here we describe the data sources and standardization procedure. All instances were converted into five-option multiple-choice questions with a single correct label, enabling binary outcome supervision for trajectory-level optimization. The microbiome component comprised MDIPID-derived mechanistic inference questions spanning microbiota–drug–disease contexts [55]. The retrieval component used fact-seeking queries adapted from Natural Questions (NQ)[146]. General biomedical questions were drawn from PubMedQA and MedQA[147, 148], and mathematical reasoning questions were sampled from DeepMath-103K [149]. To improve data quality and prevent benchmark leakage, we removed near-duplicate items, normalized entity strings (for example, gene and microbe synonyms), and excluded any examples overlapping with the evaluation benchmarks.

#### Benchmark evaluation suite

We report benchmark performance in the main text (Main Text 3.1) and summarize the evaluation suite in Extended Data Fig. 2**c**. Here we define the benchmark components and the provenance of the domain-specific tasks. For general biomedical evaluation, we used the Medicine subset of MedMCQA[58] and WMDP-Bio[59], and scored accuracy against the provided gold labels. To evaluate microbiome-focused mechanistic reasoning, we curated four domain-specific multiple-choice tasks derived from MDIPID [55]: Drug-Imp, MB-Mec, Prot-Func, and Prot-Gen. Representative examples and task templates are provided below. Across all benchmarks, we standardized entity representations and applied the same leakage-prevention filters described above.

#### Task 1: Drug–Microbe Impact (Drug-Imp)

This task tests whether the system can identify bacterial taxa that exhibit directional changes (enrichment or depletion) in response to pharmaceutical agents or dietary interventions, using curated directional effects in the gut.

##### Task example 1: Drug–Microbe Impact (Drug-Imp)

**Query:**

In a stool-based observational study evaluating the impact of a vegetarian diet on gut microbiota composition, researchers reported that this dietary pattern decreased the relative abundance of a specific bacterial taxon. Which of the following gut microbial taxa was found to be reduced in relative abundance in individuals adhering to a vegetarian diet?

A. Roseburia sp.
B. Collinsella sp.
C. Mediterraneibacter gnavus
D. Ruminococcus torques
E. unclassified Lachnospiraceae

**Answer:** B. Collinsella sp.

#### Task 2: Microbe–Protein Mechanism (MB-Mec)

To assess mechanistic reasoning, this task asks the system to pinpoint the enzymes or functional proteins that mediate a microbe’s metabolic interaction with a drug, linking organism-level phenotypes to specific molecular drivers.

##### Task example 2: Microbe–Protein Mechanism (MB-Mec)

**Query:**

In Streptococcus mitis, which specific membrane-associated enzyme is most likely responsible for the microbiome-mediated metabolic modification of daptomycin by catalyzing the transfer of phosphatidyl groups from CDP-activated alcohols to diacylglycerol during phospholipid biosynthesis, thereby altering the bacterial membrane composition and impacting daptomycin activity?

A. CDP-alcohol phosphatidyltransferase
B. Multiple sugar-binding transport ATP-binding protein msmK
C. Xylose isomerase
D. NADPH-dependent curcumin reductase
E. Cytochrome P450 monooxygenase 51A

**Answer:** A. CDP-alcohol phosphatidyltransferase

#### Task 3: Protein Functional Comprehension (Prot-Func)

This task requires identifying the precise biological function of a protein within a specified microbial species by selecting the correct activity among closely related biochemical alternatives.

##### Task example 3: Protein Functional Comprehension (Prot-Func)

**Query:**

Which of the following descriptions best characterizes the biological function of the protein “Linoleate 10-hydratase” found in Lactiplantibacillus plantarum?

A. This enzyme catalyzes the carboxylation of acetone to form acetoacetate, and it has a reduced activity on butanone, and no activity on 2-pentatone, 3-pentatone, 2-hexanone, chloroacetone, pyruvate, phosphoenolpyruvate, acetaldehyde, propionaldehyde and propylene oxide. And it requires Mg2+ and ATP.
B. This enzyme catalyzes the committed step in the biosynthesis of acidic phospholipids known by the common names phophatidylglycerols and cardiolipins.
C. This enzyme is a pyridoxal-phosphate protein providing the (R)-glutamate required for cell wall biosynthesis and converting L- or D-glutamate to D- or L-glutamate, respectively, but not other amino acids such as alanine, aspartate, and glutamine.
D. This enzyme catalyzes the reductive cleavage of azo bond in aromatic azo compounds to the corresponding amines.
E. This enzyme catalyzes the conversion of linoleic acid to 10-hydroxy-12-octadecenoic acid.

**Answer:** E. This enzyme catalyzes the conversion of linoleic acid to 10-hydroxy-12-octadecenoic acid.

#### Task 4: Protein–Gene Mapping (Prot-Gen)

This task maps functional protein descriptions to standardized gene names in the corresponding organism, testing precision in genomic grounding.

##### Task example 4: Protein–Gene Mapping (Prot-Gen)

**Query:**

In the genome of Klebsiella pneumoniae, which standardized gene is responsible for encoding the protein ‘Molybdopterin-dependent enzyme’?

A. molD
B. acxB
C. Lsa784
D. yieF
E. BSAG_01549

**Answer:** A. molD

#### Scientist preference evaluation queries

Domain experts in microbiology and immunology compared blinded, side-by-side responses from **Eubiota** and GPT-5.1. Representative evaluation queries, spanning foundational microbiome concepts and specific research mechanisms, included:

##### Representative queries used in the human evaluation study

**Microbiome–immune interactions**

- What agonist from bifidobacterium breve leads to IL-10 production by dendritic cells?
- What is the most immunogenic commensal bacteria in the human gut?

**Gut bacterial composition and pathogenicity**

- Which genus of microbiota is the most prevalent in the adult microbiome
- Why are Proteobacteria commonly associated with being pathobionts?

**Diet–microbiome interactions**

- How does the type of fat in the diet influence gut microbes through bile acids?
- Which gut microbial products may explain the link between red meat intake and inflammation?

### Supplementary Note 6: Experimental resources and protocols

**Supplementary Table 1:** Top-ranked genes across different selection strategies.

| # | Gene name | Final position | Gene ID |
| --- | --- | --- | --- |
| **Top 10 of the global full list** |  |  |  |
| 1 | *obgE* | P2-A11 | BBR_RS19450 |
| 2 | *ruvB* | P15-B3 | BBR_RS15660 |
| 3 | *dusB* | P4-C5 | BBR_RS17165 |
| 4 | *rlmH* | P16-B5 | BBR_RS19685 |
| 5 | *truA* | P15-D11 | BBR_RS18795 |
| 6 | *uvrA* | P12-D1 | BBR_RS13555 |
| 7 | *ruvA* | P15-H8 | BBR_RS15655 |
| 8 | *ychF* | P13-E7 | BBR_RS14650 |
| 9 | *uvrC* | P6-G10 | BBR_RS15515 |
| 10 | *trmB* | P6-E5 | BBR_RS18245 |
| **Top 10 of the random selection** |  |  |  |
| 1 | *lgt* | P4-F12 | BBR_RS15805 |
| 2 | *hisD* | P7-A11 | BBR_RS17055 |
| 3 | *licT* | P14-F7 | BBR_RS18695 |
| 4 | *lepA* | P4-C8 | BBR_RS15230 |
| 5 | *hpt* | P12-A6 | BBR_RS18185 |
| 6 | *purT* | P16-F9 | BBR_RS13455 |
| 7 | *trxA* | P15-A12 | BBR_RS12835 |
| 8 | *nadD* | P5-C12 | BBR_RS14085 |
| 9 | *pyrC* | P14-C6 | BBR_RS15435 |
| 10 | *rbsC* | P12-A2 | BBR_RS17765 |
| **Top 10 of the 200-gene list** |  |  |  |
| 1 | *uvrA* | P12-D1 | BBR_RS13555 |
| 2 | *recR* | P4-C11 | BBR_RS10990 |
| 3 | *lgt* | P4-F12 | BBR_RS15805 |
| 4 | *tgt* | P1-G5 | BBR_RS10715 |
| 5 | *ahpC* | P1-B10 | BBR_RS10460 |
| 6 | *dusB* | P4-C5 | BBR_RS17165 |
| 7 | *obgE* | P2-A11 | BBR_RS19450 |
| 8 | *ilvA* | P2-D3 | BBR_RS20130 |
| 9 | *hpt* | P12-A6 | BBR_RS18185 |
| 10 | *nfo* | P7-G3 | BBR_RS15825 |

**Supplementary Table 2:** Bacterial strains and respective growth medium.

| **Bacteria Strain** | **Medium** |
| --- | --- |
| *Dorea formicigenerans* ATCC 27755 | BHI + 5 µg/mL Hemin + 1 µg/mL Vitamin K3 |
| *Ruminococcus gnavus* ATCC 29149 | BHI + 5 µg/mL Hemin + 1 µg/mL Vitamin K3 |
| *Blautia obeum* ATCC 29174 | BHI + 5 µg/mL Hemin + 1 µg/mL Vitamin K3 |
| *Lactobacillus ruminis* ATCC 25644 | BHI + 5 µg/mL Hemin + 1 µg/mL Vitamin K3 |
| *Lactobacillus plantarum* ATCC 14917 | BHI + 5 µg/mL Hemin + 1 µg/mL Vitamin K3 |
| *Streptococcus thermophilus* ATCC 19258 | BHI + 5 µg/mL Hemin + 1 µg/mL Vitamin K3 |
| *Blautia hansenii* DSM 20583 | Modified Columbia medium |
| *Bifidobacterium breve* DSM 20213 | Modified Columbia medium |
| *Faecalibacterium prausnitzii* DSM 17677 | Modified Columbia medium |
| *Roseburia intestinalis* DSM 14610 | YCFAC medium |
| *Bifidobacterium breve* UCC2003 inoculum selection | YCFAC (no cysteine) + 10 µg/mL Tetracycline |
| *Bifidobacterium breve* UCC2003 growth curve experiments | YCFAC (no cysteine) |
| *Pseudomonas aeruginosa* PA14 | BHI (Aerobic/Anaerobic) |
| *Citrobacter portucalensis* HM-299 | BHI (Aerobic/Anaerobic) |
| *Klebsiella pneumoniae* NCTC 9633 | BHI (Aerobic/Anaerobic) |
| *Escherichia coli* K12 | BHI (Aerobic/Anaerobic) |
| *Bacteroides thetaiotaomicron* VPI-5482 | BHI (Anaerobic) |
| *Bacteroides ovatus* ATCC 8483 | BHI (Anaerobic) |
| *Bifidobacterium breve* DSM 20213 | BHI (Anaerobic) |

#### Laboratory list of 22 strains available for therapeutic design

*Bacteroides fragilis, Bacteroides thetaiotaomicron, Bifidobacterium breve, Lactococcus lactis, Roseburia intestinalis, Blautia hansenii, Lactobacillus ruminis, Bacteroides finegoldii, Parabacteroides johnsonii, Bacteroides vulgatus, Bilophila wadsworthia, Bacteroides stercoris, Faecalibacterium prausnitzii, Dorea longicatena, Butyricimonas virosa, Coprococcus comes, Dorea formicigenerans, Ruminococcus gnavus, Blautia obeum, Lactobacillus plantarum, Streptococcus thermophilus, Intestinimonas butyriciproducens*

#### Supplementary protocols

As part of the agentic discovery workflow, **Eubiota** generated structured experimental protocols based on experimental goals, a database of protocols provided by scientists, and standard laboratory practices. The protocols shown in this section are representative examples of experimental protocols generated by the **Eubiota** system. Sections marked gray were not used for wet lab validation.

##### Sample protocol 1: Genetic discovery

1. **High-Level Summary** This protocol outlines a comprehensive experimental plan to assess the impact of specific candidate genes on gut bacteria responses to nitric oxide (NO) exposure. The experiment involves anaerobic bacterial culture, growth curve analysis under NO stress, gene function assays, and appropriate experimental controls to determine gene-specific effects on bacterial fitness.
2. **Experiment Modules** **Module 1: Bacterial Culture** **Objective:** Anaerobically culture selected bacterial strains to prepare for growth curve analysis.

**Materials & Equipment:** Coy anaerobic chamber, 37 °C incubator, 24-deep well plate, AeroSeal™ film, multichannel pipettes, sterile tips, YCFAC or modified Columbia medium.

- **Bacterial culture preparation**
  1. Prepare and sterilize base media without cysteine. Add hemin and vitamin K_3_ supplements and pre-reduce media overnight at 37 °C.
  2. Retrieve glycerol stocks from −80 °C and inoculate 2 mL of pre-reduced medium with bacterial strains in a 24-deep well plate. Include media-only controls.
  3. Seal plates with AeroSeal™ film and incubate anaerobically at 37 °C for 48–72 h.

**Module 2: Growth Curve Analysis**

**Objective:** Measure bacterial growth under nitric oxide exposure to assess gene function.

**Materials & Equipment:** 96-well plate, Epoch 2 microplate reader, NO stock solutions, multichannel pipettes.

- **Growth curve measurement under NO stress**
  1. Prepare NO stock solutions and perform serial dilutions to achieve desired concentrations. Dilute bacterial cultures and dispense into 96-well plates.
  2. Configure the microplate reader to measure absorbance at 600 nm every 15 min with double orbital shaking at 37 °C.
  3. Run measurements for 48–72 h, export raw data to Excel, and organize for downstream analysis.

**Module 3: Gene Function Assays**

**Objective:** Assess the function of candidate genes using gene knockout or overexpression approaches.

**Materials & Equipment:** Transposon library, PCR reagents, qPCR instrumentation.

- **Gene perturbation and validation**
  1. Use transposon libraries to find candidate genes in bacterial mutants.
  2. Validate gene disruptions or overexpression using PCR and quantitative PCR.
  3. Compare growth curves of genetically modified strains under NO exposure to wild-type controls.

**Module 4: Controls and Data Analysis**

**Objective:** Implement appropriate controls and analyze data to ensure experimental validity.

**Materials & Equipment:** Excel, statistical analysis software.

- **Controls and analysis**
  1. Include media-only and untreated bacterial controls in all experiments.
  2. Analyze growth curves using statistical software, comparing NO-treated and control groups following established methods *(Atolia et al*., *mBio, 2020)*.

##### Sample protocol 2: Microbial consortium design for gut diseases

1. **High-Level Summary** This protocol outlines the steps to establish a DSS-induced colitis model in mice, administer probiotic communities, and quantify inflammatory markers including lipocalin-2. The experimental workflow includes defined timelines for DSS exposure, probiotic gavage, sample collection, and tissue processing. The protocol is organized into modular sections covering colitis induction, probiotic administration, and inflammation marker measurement.
2. **Experiment Modules**

**Module 1: DSS-Induced Colitis Model Setup**

**Objective:** Induce colitis in mice using DSS to study intestinal inflammation.

**Materials:** DSS (2.5% in drinking water), C57BL/6 mice, animal balance, gavage needles, syringes, microcentrifuge tubes.

- **DSS colitis induction and monitoring**
  1. **Day 0:** Prepare DSS solution by dissolving DSS powder in autoclaved drinking water. Provide DSS water *ad libitum* for 5–7 days.
  2. **Daily monitoring:** Record body weight, stool consistency, and fecal blood using a Hemoccult test.
  3. **Euthanasia and tissue collection:** Perform euthanasia, collect colon tissue and cecal contents, and measure colon length.

**Module 2: Probiotic Community Administration**

**Objective:** Administer probiotic communities to evaluate their effects on DSS-induced colitis.

**Materials:** Anaerobic chamber, pre-reduced culture medium, selected bacterial strains.

- **Probiotic gavage regimen**
  1. **Day** −2: Gavage mice with 200 *µ*L of the synthetic microbial community daily for two days prior to DSS administration.
  2. **Day 0–end point:** Continue gavage every two days throughout DSS treatment and the recovery phase.

**Module 3: Inflammation Marker Measurement**

**Objective:** Quantify inflammatory lipocalin-2 in collected samples.

**Materials:** ELISA kits for lipocalin-2, microplates, plate reader.

- **Cytokine quantification by ELISA**
  1. **Sample collection:** Collect blood and tissue samples at designated time points during and after DSS treatment.
  2. **ELISA setup:** Coat microplates with capture antibodies, block nonspecific binding sites, and add samples.
  3. **Detection:** Add detection antibodies, substrate, and stop solution. Measure absorbance at 450 nm.

##### Sample protocol 3: Pathogen-biased antibiotic cocktail design

1. **High-Level Summary** This protocol outlines a comprehensive experimental plan to evaluate the efficacy of three antibiotics—Piperacillin, Tazobactam, and Amikacin—against pathogenic bacteria, including *Pseudomonas aeruginosa, Citrobacter portucalensis*, and *Klebsiella pneumoniae*. In parallel, the protocol assesses the collateral impact of these antibiotics on beneficial gut bacteria, including *Escherichia coli, Bacteroides* spp., and *Bifidobacterium* spp. The workflow is organized into modular sections covering bacterial culture, antibiotic efficacy testing, impact assessment on gut bacteria, and data analysis.
2. **Experiment Modules**

**Module 1: Bacterial Culture Preparation**

**Objective:** Culture pathogenic and beneficial bacterial strains under appropriate aerobic and anaerobic conditions.

- **Bacterial culture setup**
  1. **Anaerobic culture:** Use PROT-1 to culture *Bacteroides* spp. and *Bifidobacterium* spp. anaerobically in a Coy chamber at 37 °C for 48–72 h.
  2. **Aerobic culture:** Use PROT-1 to culture *Pseudomonas aeruginosa, Citrobacter portucalensis, Klebsiella pneumoniae*, and *Escherichia coli* aerobically in a shaking incubator at 37 °C with agitation at 200–250 rpm.

**Module 2: Antibiotic Efficacy Testing**

**Objective:** Assess the efficacy of antibiotics against pathogenic bacterial strains.

- **Antibiotic susceptibility assays**
  1. Perform spot dilution assays on agar plates containing Piperacillin, Tazobactam, Amikacin at varying concentrations following PROT-54.
  2. Quantify colony-forming units (CFUs) to determine bacterial survival rates following antibiotic exposure.

**Module 3: Impact on Gut Bacteria**

**Objective:** Evaluate the impact of antibiotic treatment on beneficial gut bacterial species.

- **Gut microbiota impact assessment**
  1. Prepare gut bacterial cultures using the same aerobic or anaerobic conditions described in Module 1.
  2. Expose cultures to the same antibiotic concentrations used in Module 2.
  3. Assess bacterial viability using the LIVE/DEAD BacLight bacterial viability assay

**Module 4: Data Analysis and Interpretation**

**Objective:** Analyze and interpret data from antibiotic efficacy and gut microbiota impact studies.

- **Data analysis and interpretation**
  1. Compile data from CFU quantification and bacterial viability assays.
  2. Perform statistical analyses using software such as GraphPad Prism, including ANOVA and appropriate post hoc tests.
  3. Compare antibiotic efficacy against pathogenic bacteria with corresponding effects on beneficial gut bacteria.

##### Sample protocol 4: Anti-inflammatory molecule discovery

1. **High-Level Summary** This protocol outlines the steps to test 5 candidate metabolites for NF-*κ*B inhibition in macrophages using flow cytometry.The experiment involves macrophage culture, treatment with metabolites, LPS stimulation, and measurement of NF-*κ*B activation. The protocol is divided into modules, each detailing specific experimental procedures, some mapped to provided protocols and others derived from standard laboratory practices.
2. **Experiment Modules**

**Module 1: Macrophage Culture and Seeding**

- **Day −2: Seed macrophage cells**
  1. Inspect cell density and refresh media if necessary.
  2. Harvest cells by removing old media, washing, and resuspending in fresh media.
  3. Count cells using a hemocytometer and calculate seeding density.
  4. Seed 75,000 cells per well in a 96-well plate.

**Module 2: Metabolite Preparation**

- **Methanol extraction of metabolites**
  1. Add 100 *µ*L of sample to each well of a 96 deep-well plate.
  2. Add 1,000 *µ*L of 100% MeOH and incubate at −20 °C for 30–60 min.
  3. Centrifuge at 5,000 ×*g* for 5 min, transfer supernatant, and dry.
  4. Resuspend in 200 *µ*L sterile water and sterile-filter.

**Module 3: Macrophage Treatment with Metabolites**

- **Day −1: Treat macrophages with metabolites**
  1. Thaw metabolite samples and add to macrophage cultures at a final dilution of 1:200.
  2. Mix plates gently and incubate under standard culture conditions.

**Module 4: LPS Stimulation**

- **Stimulate NF-***κ***B activation**
  1. Prepare a 10-fold diluted LPS solution.
  2. Add 2 *µ*L of diluted LPS to each well after 2 h of metabolite treatment.

**Module 5: Flow Cytometry for NF-***κ***B Activation**

- **Measure NF-***κ***B activation**
  1. Harvest cells following LPS stimulation.
  2. Stain cells with antibodies specific for NF-*κ*B activation markers.
  3. Analyze samples by flow cytometry to quantify NF-*κ*B inhibition.

### Supplementary Note 7: Additional experimental analysis

**Supplementary Figure 4:**
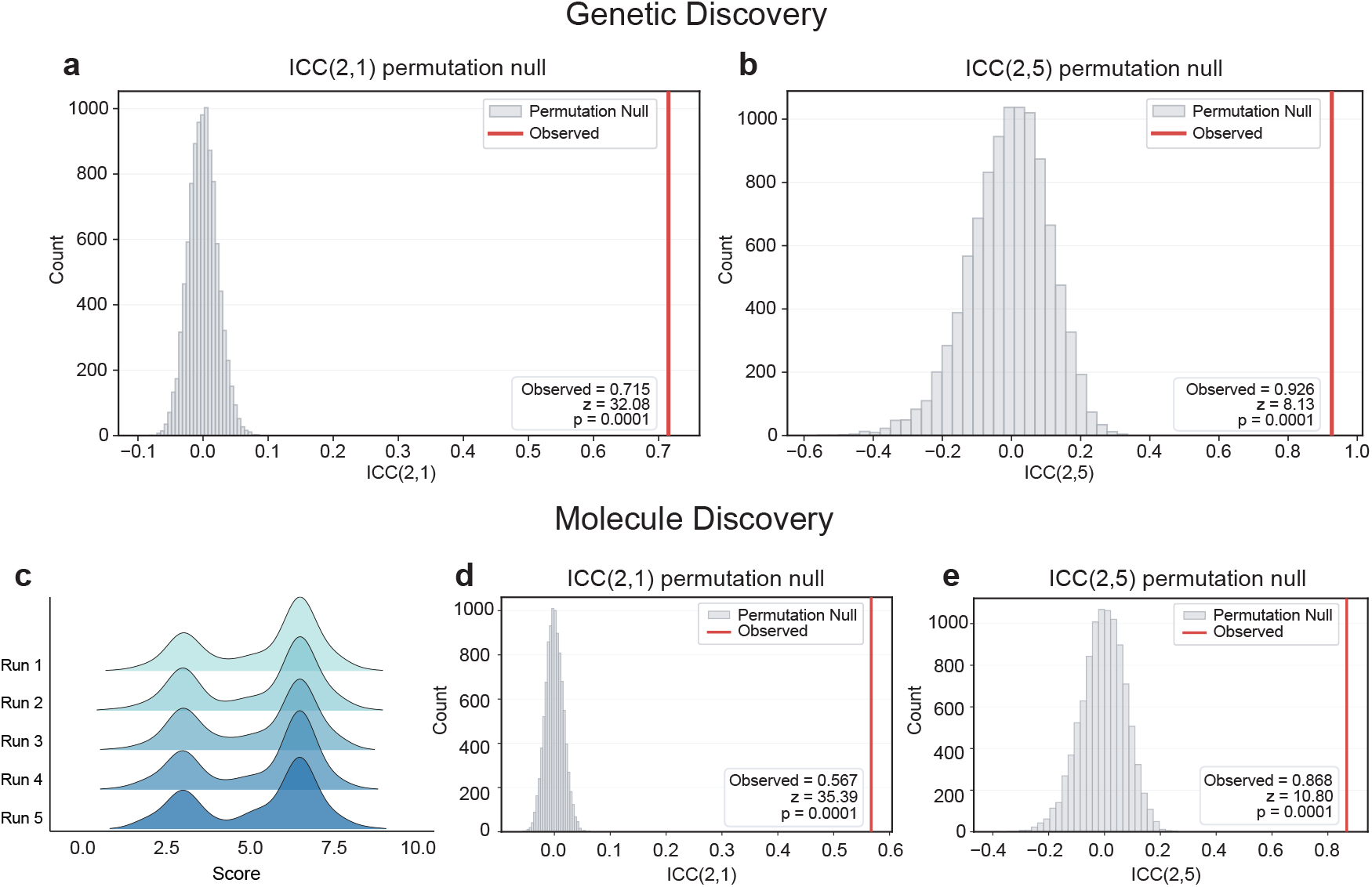
Statistical stability and reproducibility of Eubiota reasoning across independent runs. **a, b**, ICC(2,1) and ICC(2,5) permutation tests of the genetic discovery test. Gene scores were randomly assigned to gene labels 10,000 times to construct a null distribution. Observed ICC values are marked by the red line, with empirical *P* < 0.0001. **c**, Ridgeline plot showing metabolite score distributions across five runs of the anti-inflammatory metabolite discovery task. No significant shifts in central tendency or dispersion were observed. **d, e**, ICC(2,1) and ICC(2,5) permutation tests of the molecule discovery test. Gene scores were randomly assigned to metabolite labels 10,000 times to construct a null distribution. Observed ICC values are marked by the red line, with empirical *P* < 0.0001. *P* values were calculated as the proportion of permuted ICC values that were greater than or equal to the observed ICC.

## Footnotes

1 https://platform.openai.com/docs/models/text-embedding-3-large.

2 https://ai.google.dev/gemini-api/docs/google-search.

3 https://docs.perplexity.ai.

4 https://www.wikipedia.org.

5 https://www.ncbi.nlm.nih.gov/home/develop/api.

6 https://www.kegg.jp/kegg/rest.

7 https://mdipid.idrblab.net.

